# Prioritization of osteoporosis-associated GWAS SNPs using epigenomics and transcriptomics

**DOI:** 10.1101/2020.06.18.160309

**Authors:** Xiao Zhang, Hong-Wen Deng, Hui Shen, Melanie Ehrlich

**Affiliations:** Department of Biostatistics and Data Science, School of Public Health and Tropical Medicine, Tulane University, New Orleans, LA, USA; Tulane Center for Biomedical Informatics and Genomics, Division of Biomedical Informatics and Genomics, Deming Department of Medicine, School of Medicine, Tulane University, New Orleans, LA, USA; Tulane Cancer Center and Hayward Genetics Center, Tulane University, New Orleans, LA, USA

**Keywords:** osteoporosis, osteoblasts, epigenetics, molecular pathways – remodeling, Wnt/β-catenin

## Abstract

Genetic risk factors for osteoporosis, a prevalent disease associated with aging, have been examined in many genome-wide association studies (GWAS). A major challenge is to prioritize transcription-regulatory GWAS-derived variants that are likely to be functional. Given the critical role of epigenetics in gene regulation, we have used an unusual epigenetics- and transcription-based approach to identify credible regulatory SNPs relevant to osteoporosis from 38 reported BMD GWAS. Using Roadmap databases, we prioritized SNPs based upon their overlap with strong enhancer or promoter chromatin preferentially in osteoblasts relative to 11 heterologous cell culture types. The selected SNPs also had to overlap open chromatin (DNaseI-hypersensitive sites) and DNA sequences predicted to bind to osteoblast-relevant transcription factors in an allele-specific manner. From >50,000 GWAS-derived SNPs, we identified 16 novel and credible regulatory SNPs (Tier-1 SNPs) for osteoporosis risk. Their associated genes, *BICC1, LGR4, DAAM2, NPR3*, or *HMGA2*, are involved in osteoblastogenesis or bone homeostasis and regulate cell signaling or enhancer function. Four of them are preferentially expressed in osteoblasts. *BICC1, LGR4*, and *DAAM2* play important roles in canonical Wnt signaling, a pathway critical to bone formation and repair. The transcription factors that are predicted to bind to the Tier-1 SNP-containing DNA sequences also have bone-related functions. For the seven Tier-1 SNPs near the 5’ end of *BICC1*, examination of eQTL overlap and the distribution of BMD-increasing alleles suggests that at least one SNP in each of two clusters contributes to inherited osteoporosis risk. Our study not only illustrates a method that can be used to identify novel BMD-related causal regulatory SNPs for future study, but also reveals evidence that some of the Tier-1 SNPs exert their effects on BMD risk indirectly through little-studied noncoding RNA genes, which in turn may control the nearby bone-related protein-encoding gene.

## Introduction

Osteoporosis, which affects about 200 million people worldwide, involves a loss of bone mass, strength, and normal microarchitecture.^(1)^ Both osteoporosis and bone fracture increase greatly with age.^(2)^ Bone mineral density (BMD) of the hip, spine, limb, or total body is the quantitative trait used most frequently to detect or predict osteoporosis.^(1, 3)^ In the clinic, BMD is usually determined by dual-energy x-ray absorptiometry (DXA). In genome-wide association studies (GWAS), the simpler-to-perform ultrasound-measured heel bone density (estimated BMD, eBMD) is often used. Like DXA-determined BMD, eBMD has a high heritability (50 - 80%) and can predict bone fracture after correcting for the effects of height, weight, age, and sex.^(4)^ Even low-frequency variants (minor allele frequency of 1 - 5%) or rare variants may contribute to decreased BMD and increased bone fracture.^(5)^

From DXA or eBMD GWAS (collectively referred to here as BMD GWAS), hundreds of genes have been identified as having nearby or overlapping genetic variants significantly associated with osteoporosis risk.^(1, 4, 6)^ Most BMD or other trait-associated variants are located in non-coding regions of the genome.^(1, 7)^ Identifying causal osteoporosis-risk genes and their associated variants, usually SNPs, is challenging for variants that are not missense coding variants. The GWAS-derived SNP exhibiting the smallest p-value for association with the studied trait at a given locus (index SNP) is often linked to very many proxy SNPs solely because of linkage disequilibrium (LD) rather than due to biological relevance. Recently, whole-genome profiles of epigenetics features, such as overlap with open chromatin (e.g., DNaseI hypersensitive sites, DHS), have been used to help discriminate likely causal BMD regulatory variants from bystander variants in high LD with them.^(5, 6, 8-10)^ Moreover, GWAS SNPs are generally also enriched in enhancer or promoter chromatin.^(7, 10)^ The importance of considering such epigenetic features at BMD GWAS-derived SNPs is also evidenced by the finding that specific epigenetic changes play a key role in bone formation and remodeling.^(11, 12)^

Using epigenetics to help prioritize BMD GWAS variants requires choosing the cell types for study. Bone has one of the most complex developmental and repair pathways due in part to its rigidity, strength, highly dynamic nature, and multiple structural and physiological roles.^(13)^ Osteoblasts (ostb) play a central role in both intramembranous ossification and in the more widespread and complicated endochondral ossification pathway for bone formation based upon chondrocytes (chond).^(11)^ Surprisingly, there is evidence for hypertrophic chond transdifferentiating to ostb, which is consistent with the considerable overlap of transcription profiles of these two morphologically dissimilar bone cell types.^(14, 15)^ Ostb and chond are derived from a heterogeneous mixture of bone-derived stem or stem-like cells commonly referred to as mesenchymal stem cells (MSC).^(13)^ Bone-destroying osteoclasts, which arise from the monocyte blood lineage, contribute to osteoporosis when bone resorption is excessive. Remodeling of cortical bone occurs through complex multicellular units containing ostb and osteoclasts, which modulate each other’s function and are often responsive to some of the same growth factors and cytokines, although sometimes with different responses.^(2, 16)^ Osteocytes (terminally differentiated ostb), which constitute most of the cells in cortical bone, coordinate the differentiation and activity of ostb and osteoclasts in response to mechanical load or injury.^(17)^

Because of the centrality of ostb to normal bone homeostasis and the availability of epigenomic and transcriptomic profiles for comparing human cell cultures of ostb, chond, MSC, and heterologous primary cell cultures, we focused on examining the epigenetics of BMD GWAS-derived SNPs in ostb and secondarily, in chond and MSC. In a recent study, we used epigenetics to prioritize best-candidate causal variants for BMD GWAS SNPs and obesity GWAS SNPs at a single gene, *TBX15*, which encodes a limb development-associated transcription factor (TF).^(18)^ In the present study, we take the unusual approach of looking genome-wide not only for SNPs overlaying enhancer or promoter chromatin in ostb but also for such SNPs that displayed these types of regulatory chromatin preferentially in ostb relative to many types of cell cultures not directly related to bone. Using epigenomics, transcriptomics, gene ontology (GO) analysis, and TF binding site (TFBS) prediction in a analysis of whole-genome data from 38 BMD GWAS, we identified 16 high-priority candidates for causal regulatory SNPs associated with five bone-relevant genes (*BICC1, NPR3, LGR4, HMGA2*, and *DAAM2*). None of these SNPs was previously described as associated with osteoporosis risk.

## Materials and Methods BMD GWAS-derived SNPs

Index SNPs associated with BMD (*p* < 5 × 10^−8^) were retrieved from 38 studies in the NHGRI-EBI GWAS Catalog^(19)^ (downloaded October 2019; Supplemental Table S1). We expanded the index SNPs to a set of proxy SNPs (*r*^2^ ≥ 0.8, EUR from the 1000 Genome Project Phase 3^(20)^) by using PLINK v1.9^(21)^ (https://www.cog-genomics.org/plink/1.9/) and/or LDlink v3.9^(22)^ (https://ldlink.nci.nih.gov/). For the five best candidate genes, we also extracted imputed SNPs (*p* < 5 × 10^−8^) for total-body DXA BMD^(3)^ and (*p* < 6.6 × 10^−9^) for eBMD^(6)^ to look for additional Tier-1 SNPs.

### Transcriptomics

For determining preferential expression of genes in ostb we obtained RPKM (reads per kilobase million) for ostb and 11 heterologous cell cultures (Supplemental Table S2), not including MSC or chond, from the ENCODE RNA-seq database (Supplemental Methods; http://genome.ucsc.edu). Preferential expression was defined as the ratio of ostb RPKM/median non-ostb RPKM > 5 and with a minimum of RPKM > 1 for ostb. The differentiation status of the commercially obtained ostb that had been used to generate the ENCODE RNA-seq data as well as for the epigenomic data (see below) is unclear. However, expression and chromatin segmentation profiles for *SP7, RUNX2, SPP1, IBSP, TNFRSF11B, BGLAP*, and *ALPL* indicate selective transcription of middle- and late ostb differentiation marker genes by ostb samples and chond-specific expression of *SOX5* in chond samples (Supplemental Figs. S1 and S2). For some genes, we also examined tissue expression profiles (median transcripts per million, TPM, and tissue expression QTL, eQTL) from the GTEx database^(23)^ (https://www.gtexportal.org/) and mouse expression microarray profiles from BioGPS^(24)^ (http://biogps.org/).

### Epigenomics

Chromatin state segmentation data for the 15 examined cell culture types (Supplemental Methods and Table S3) were obtained from the 18-state Roadmap model with strong promoter chromatin as state 1 and strong enhancer chromatin as states 3, 8 or 9 (Roadmap^(7)^; http://genome.ucsc.edu). In figures for clarity in depicting these data, we slightly modified the color-coding of the chromatin state segmentation as indicated. DHS were defined as narrowPeaks from DNase-seq (Roadmap^(7)^; http://genome.ucsc.edu).

### Predictions of allele-specific transcription factor binding sites and analyses of gene functional terms

Predictions of allele-specific TFBS were made using a 21-base sequence centered around the SNP of interest for comparison to the TRANSFAC v2019.3 database (Supplemental Methods). We checked that each TF for a matching TFBS had appreciable expression in ostb (RPKM ≥ 0.8). In addition, we used manual curation to retain only those TRANSFAC TFBS predictions for which all of the conserved positions (5 – 10) in the PWM (position weight matrix) had exact matches to those in the SNP-containing sequence and no more than one base in a partly conserved position had only a partial match (at least 20% as good as the best PWM match). For determining that a TFBS overlapping a SNP is likely to be allele-specific in its TF binding, we required that the disfavored allele had a position probability that was at least 5-fold lower than that of the favored allele in the PWM matrix. DAVID v6.7^(25)^ (https://david-d.ncifcrf.gov/) was used for functional classification of reference genes and GREAT v4.0.4^(26)^ (http://great.stanford.edu/) for prioritization of genes linked to our epigenetically-selected GWAS SNP subset.

## Results

### Prioritization of 16 SNPs as candidate causal regulatory SNPs (Tier-1 SNPs) from 38 BMD GWAS

To identify high-priority candidates for causative regulatory variants from BMD GWAS, we obtained index SNPs from 38 eBMD or DXA BMD studies (Fig. 1; Supplemental Table S1) and expanded them (LD threshold of *r*^2^ ≥ 0.8) using the European ancestry (EUR) population because it was the predominant one studied. As expected, the GO term with the most enrichment among the 2,157 reference genes associated with the BMD GWAS index SNPs was skeletal system development (Supplemental Table S4). In accord with other analyses,^(1, 7)^ only a small percentage (2%, 1,139 SNPs) of the 57,235 BMD-related index/proxy SNPs mapped to coding regions, and they were associated with 29% of the reference genes (Supplemental Table S3).

**Figure 1.**
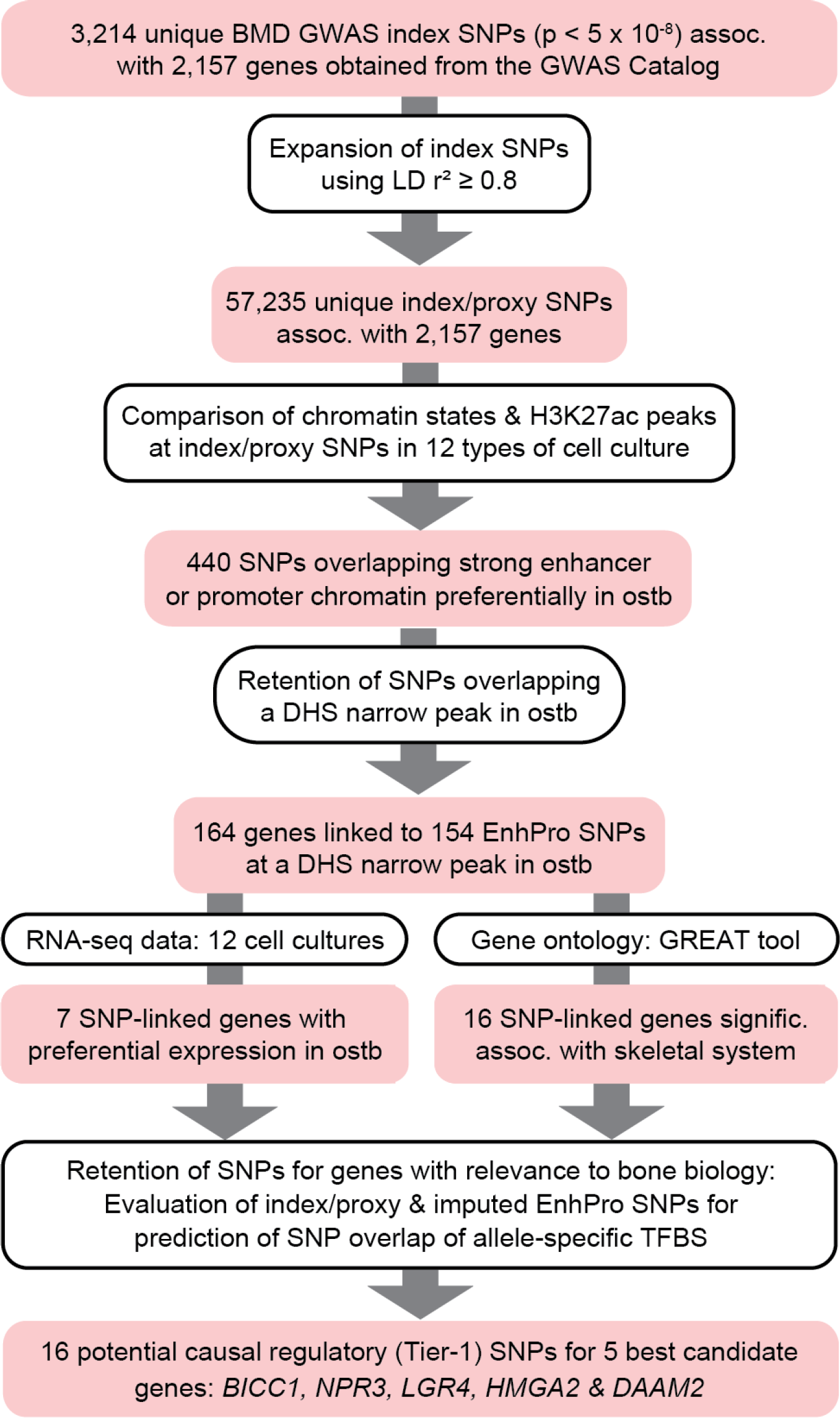
Workflow for prioritizing 16 potential causal regulatory (Tier-1) SNPs from 38 BMD GWAS. Summary of prioritization of plausible regulatory BMD SNPs. Assoc, associated; H3K27ac, histone H3 lysine-27 acetylation (an epigenetic mark of active enhancers or promoters); ostb, osteoblasts; TFBS, transcription factor binding site; DHS, DNaseI hypersensitive site; GREAT tool, http://great.stanford.edu/public/html/.

In order to prioritize GWAS SNPs for the best causative regulatory candidates, we first looked for SNPs that preferentially overlap strong transcription-regulatory enhancer or promoter chromatin in ostb. These chromatin states had been determined from genome-wide profiles of histone H3 lysine-4 trimethylation (H3K4me3) and H3 K27 acetylation (H3K27ac) for promoter chromatin and H3K4me1 and H3K27ac for enhancer chromatin.^(7)^ We found that only ∼0.8% (440) of the 57,235 index/proxy SNPs were preferentially associated with these types of chromatin in ostb and, in addition, overlapped a narrow peak of H3K27ac preferentially in this cell type (Fig. 1; Supplemental Table S3). Preferential overlap of positive-regulatory chromatin in ostb was defined as such overlap in ostb but in no more than three of the following 12 cell cultures from the Roadmap database^(7)^ (Supplemental Table S3): skin fibroblasts (fib), adult or fetal lung fib, foreskin or dermal melanocytes, keratinocytes, astrocytes, umbilical cord endothelial cells, myoblasts (myob), mammary epithelial cells (HMEC), embryonic stem cells, and a lymphoblastoid cell line. Subsequent filtering for overlap of open chromatin (DHS) in ostb gave 154 SNPs, which are referred to as EnhPro SNPs (Supplemental Table S3). EnhPro SNPs often also overlapped strong enhancer or promoter chromatin in bone marrow-derived MSC (74%) or in chond (79%) but not in monocytes (16%).

In order to identify EnhPro SNPs most likely to contribute to inherited osteoporosis risk, we used two alternate methods to evaluate their associated genes (Fig. 1). First, we identified seven EnhPro SNP-associated genes, *RUNX2, TBX15, ADAM12, NPR3, BICC1, LGR4*, and *SPECC1*, that were preferentially expressed in ostb (> 5 times more RPKM in ostb than in the median of 11 heterologous cell types). All but *SPECC1* have known bone-related functions.^(11, 18, 27-30)^ Because we recently analyzed EnhPro SNPs for *TBX15*,^(18)^ we eliminated it as well as *SPECC1* from further consideration. Alternatively, we identified seven additional EnhPro SNP-associated genes, *HMGA2, BMP5, DLX6, SIX1, SMAD3, TRPS1*, and *WNT7B*, that were strongly enriched in bone biology-related Gene Ontology (GO) terms, had literature references for their skeletal association, and displayed appreciable expression in ostb, chond, or MSC (Supplemental Table S5). The EnhPro SNPs from these prioritized genes were examined for predicted overlapping allele-specific TFBS using the TRANSFAC database and stringent hand curation. We found that four of these genes, *BICC1, NPR3, LGR4*, and *HMGA2*, had strong predictions of allele-specific TFBS overlapping at least one of their EnhPro SNPs and refer to them as Tier-1 SNPs. The fifth and last Tier-1 SNP-associated gene, *DAAM2*, was found using a slightly relaxed prioritization procedure for identification of EnhPro SNPs, as described below. In addition to the 13 index/proxy-derived Tier-1 SNPs, we found three more Tier-1 SNPs by examining imputed total body-BMD^(3)^ and eBMD^(6)^ SNPs associated with these five prioritized genes (Table 1).

**Table 1.** Sixteen best candidates for transcription-regulatory BMD-causal variants (Tier-1 SNPs) in five bone-related genes^a^.

| Gene | Tier-1 SNP for BMD association | Index/proxy (I/P) or imputed (Imp) SNP | Distance of SNP to gene TSS (kb) | Alleles (Ref/Alt) | BMD-increasing allele (Freq EUR)<sup>b</sup> | Chromatin state<sup>c</sup> at the SNP in ostb | Predicted TF binding<sup>d</sup> |  |
| --- | --- | --- | --- | --- | --- | --- | --- | --- |
|  |  |  |  |  |  |  | Ref allele | Alt allele |
| BICC1 | rs112597538 | I/P | -0.1 | T/C | Unknown (0.50) | Str prom | SREBP1/2 | None |
|  | rs1896245 | I/P & Imp | 3.2 | T/G | Alt (0.50) | Str enh/prom | SATB1 | None |
|  | rs1896243 | I/P & Imp | 3.5 | C/T | Alt (0.50) | Str enh | None | TCF3 |
|  | rs1658429 | I/P & Imp | 57.6 | T/C | Ref (0.50) | Str enh | None | AP-1 |
|  | rs11006188 | Imp | 60.6 | G/C | Ref (0.60) | Str enh | SMAD, GLIS3 | NR2F6 |
|  | rs1982173 | Imp | 61.2 | G/T | Ref (0.60) | Str enh | RBPJ | None |
|  | rs4948523 | I/P & Imp | 66.3 | A/C | Ref (0.49) | Str enh | PRDX2 | None |
| NPR3 | rs1173771 | I/P | 104.3 | A/G | Alt (0.58) | Str enh | SATB1 | None |
|  | rs7733331 | I/P | 118.1 | T/C | Alt (0.58) | Str enh | GTF2I | None |
| LGR4 | rs10835153 | I/P & Imp | -181 | A/T | Ref (0.41) | Str enh | None | BPTF |
| HMGA2 | rs80019710 | Imp | -231.1 | C/T | Alt (0.05) | Str enh | None | HMGA1 |
|  | rs17101510 | I/P & Imp | -231 | C/T | Alt (0.02) | Str enh | None | YY1 |
|  | rs12296417 | I/P & Imp | -230.5 | T/A | Alt (0.02) | Str enh | None | SATB1, ZBTB20 |
| DAAM2 | rs2504105 | I/P & Imp | 62.4 | A/G | Ref (0.40) | Str enh/prom | SMAD | None |
|  | rs2504104 | I/P & Imp | 62.4 | A/G | Ref (0.40) | Str enh/prom | None | RUNX2/1, CBFβ |
|  | rs2504103 | I/P, Imp | 62.5 | C/G | Ref (0.40) | Str enh/prom | None | NFIC |

<sup>a</sup>16 Tier-1 SNPs (prioritized candidates for regulatory osteoporosis-risk SNPs) and their 5 associated genes determined from 38 BMD GWAS. TSS, transcription start site; ostb, osteoblasts; TF, transcription factor; I/P, index or proxy SNP; Imp, imputed SNP; Ref, reference allele; Alt, alternate allele; Str enh, strong enhancer chromatin; Str prom, strong promoter chromatin
<sup>b</sup>The BMD-increasing alleles and their frequency in the European population
<sup>c</sup>From Roadmap 18-State/Auxiliary Hidden Markov Model, States 1, 3, 8 or 9
<sup>d</sup>Predictions of allele-specific TF binding sites by hand curation of data from the TRANSFAC database and verification of TF expression in ostb

### *BICC1* has seven BMD-risk Tier-1 SNPs near the 5’ end of the gene

Among the five prioritized genes, *BICC1* (BicC Family RNA Binding Protein 1) had the most Tier-1 SNPs (seven; Table 1). *BICC1* encodes a translation-regulatory protein that is implicated in osteoblastogenesis, plays critical roles in several signal transduction pathways, and is required for development and homeostasis of various organs including bone.^(27, 31, 32)^ It is the only gene in the surrounding 1-Mb region that is preferentially and appreciably expressed in ostb and is expressed more strongly in skin fib than in a variety of tissues although more weakly in skin fib than in ostb (Supplemental Figs. S3 and S4). Mouse expression microarray profiles indicate that there is much transcription of *Bicc1* in ostb and negligible expression in osteoclasts, as was the case for all the other Tier-1 SNP associated genes (Supplemental Figs. S5A and B and S6). As positive controls, we checked that osteoclast markers *Ctsk*, and *Tnfrsf11a/Rank* exhibit the expected osteoclast-associated expression on these microarrays (Supplemental Fig. S5C and D).

The seven *BICC1* Tier-1 SNPs were in two distinct clusters starting from 0.1 kb upstream of the *BICC1* transcription start site (TSS – 0.1 kb; rs112597538, rs1896245, and rs1896243) and within the central region of the 107-kb intron 1 (rs1658429, rs11006188, rs1982173, and rs4948523; Fig. 2A and Supplemental Fig. S3E). They were embedded in active promoter chromatin, mixed promoter/enhancer chromatin, or strong enhancer chromatin overlapping DHS in ostb, chond, and MSC (Fig. 2C - E), all of which display high expression of *BICC1* (Supplemental Fig. S3). Rs112597538 overlapped a constitutive nucleosome-depleted region seen as a ∼0.17-kb hole in the H3K27ac signal surrounding this SNP (triangle, Fig. 2D), which is indicative of strong nucleosome phasing due to TF binding. All seven Tier-1 SNPs are in moderate-to-high LD with each other (*r*^2^ = 0.65 to 1, EUR; Supplemental Fig. S7). Rs112597538, rs1896245, rs1658429 and rs1982173 are located in sequences that are evolutionarily conserved among mammals (Fig. 2B), a trait enriched in non-coding transcription regulatory regions.

**Figure 2.**
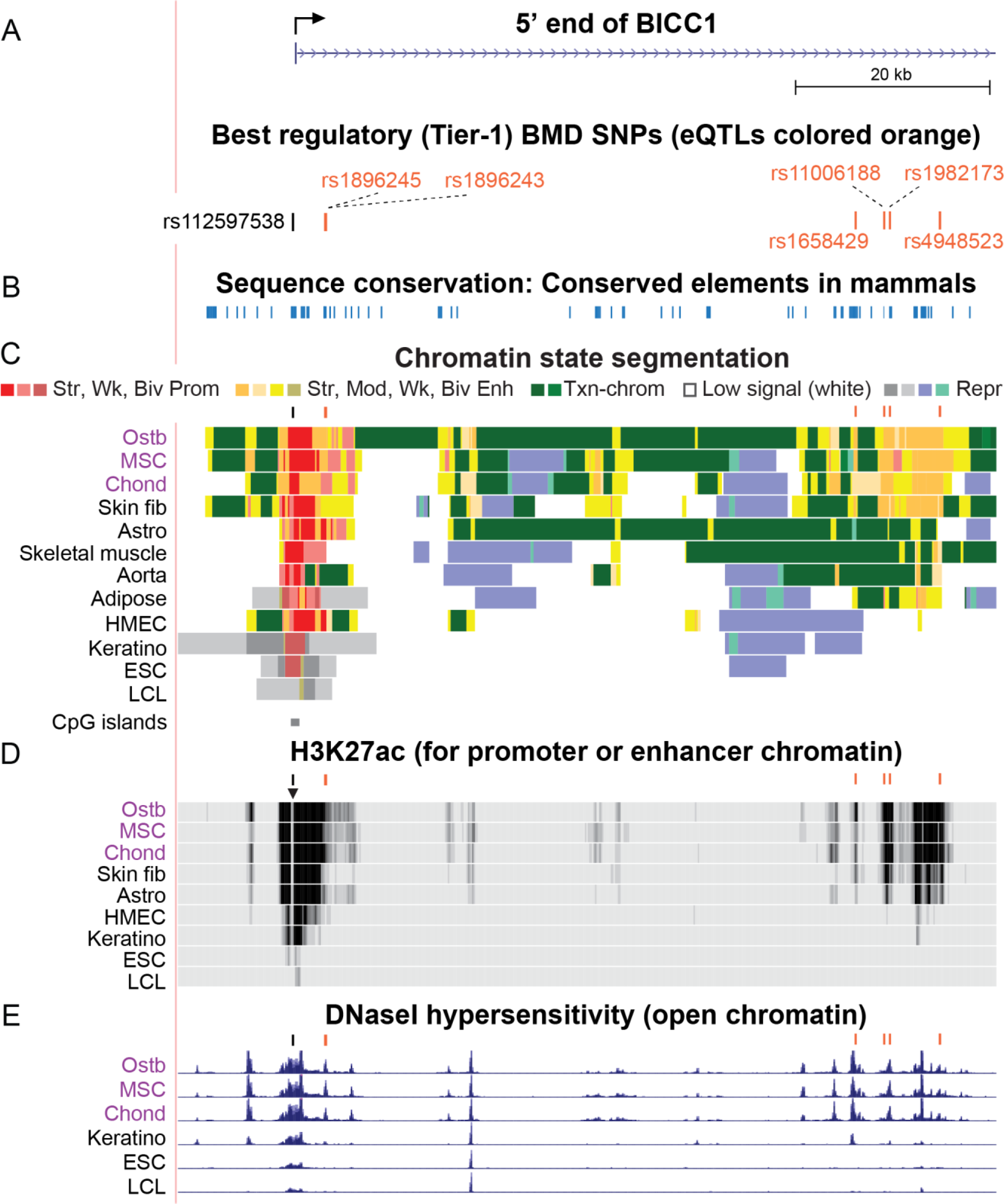
*BICC1*, which encodes an RNA-binding protein involved in regulating Wnt signaling, has two clusters of Tier-1 SNPs near its 5’ end. *(A)* An 84-kb region around the *BICC1* TSS (chr10:60,260,940-60,345,012) with seven Tier-1 SNPs near the TSS (broken arrow) or downstream in intron 1. *(B)* DNA sequence conservation among placental mammals. *(C)* Roadmap-derived chromatin state segmentation: strong (Str), moderate (Mod), weak (Wk), or bivalent (Biv; poised) promoter (Prom) or enhancer (Enh) chromatin; repressed (Repr) chromatin; or chromatin with the H3K36me3 mark of actively transcribed regions (Txn-chrom); CpG islands, CpG-rich regions. *(D)* H327ac enrichment profiles. (E) Profiles of open chromatin (DHS). Short bars above tracks in panels *C, D*, and *E*, positions of Tier-1 SNPs. All tracks were visualized in the UCSC Genome Browser (hg19) and are aligned in this figure and Figs. 3-6. Ostb, osteoblasts; MSC, mesenchymal stem cells; chond, chondrocytes; fib, fibroblasts; astro, astrocytes; HMEC, mammary epithelial cells; NHEK, foreskin keratinocytes; LCL, lymphoblastoid cell line.

Six of the *BICC1* Tier-1 SNPs were eQTLs for SkM and tibial artery (*p* = 1 × 10^−11^ to 8 × 10^−16^; no data were available for rs112597538 in the GTEx database ^(23)^). The eQTLs that were in the proximal region of *BICC1* had positive effect sizes while those in the downstream intron-1 region had negative effect sizes for many tissues including skeletal muscle and aorta (*p* = 2 × 10^−4^ to 3 × 10^−8^).^(23)^ Chromatin state profiles make the strongest case for rs112597538 at TSS - 0.1 kb and rs4948523 at TSS + 66 kb (intron 1) driving the eQTL data because these SNPs overlapped promoter or enhancer chromatin in skeletal muscle and aorta, the tissues displaying the *BICC1* eQTLs (Fig. 2C). Other Tier-1 SNPs in *BICC1* might also influence expression in ostb given the large differences in enhancer chromatin profiles of ostb from those of skeletal muscle or aorta and the lack of available eQTL profiles for ostb.

Importantly, the grouping of *BICC1* Tier-1 SNPs according to the directionality of their eQTL is consistent with their grouping by BMD-increasing allele and with their locations in *BICC1* (Table 1; Fig. 2). Further support for the biological relevance of several or more of these Tier-1 SNPs to osteoporosis is that TFs predicted to bind in an allele-specific manner to all the *BICC1* Tier-1 SNPs were related to the skeletal system (Supplemental Table S6). For example, the Reference genome (Ref) allele, and not the alternate (Alt) allele, of rs112597538 is predicted to bind to SREBP1 and SREBP2 (Table 1), which are implicated in bone biology (Supplemental Table S6).

### *NPR3* has two intergenic BMD-risk Tier-1 SNPs upstream of a novel lincRNA gene

There were two Tier-1 SNPs linked to the natriuretic peptide receptor 3 gene (*NPR3*), which encodes a receptor that facilitates clearance of natriuretic peptides to regulate cell signaling^(33)^ and is implicated in abnormal bone growth phenotypes in humans and mice.^(28)^ *NPR3* is moderately to highly expressed in ostb, chond, MSC, lung fib, myob, and skin fib and shows low or negligible expression in the other examined cell cultures (Fig. 3E). Enhancer chromatin profiles paralleled the expression data (Fig. 3C and D). Among various tissues, its highest expression is in aorta (Supplemental Fig. S4B). None of the nearby genes within its 1-Mb neighborhood has a transcription profile like that of *NPR3* (Supplemental Table S7).

**Figure 3.**
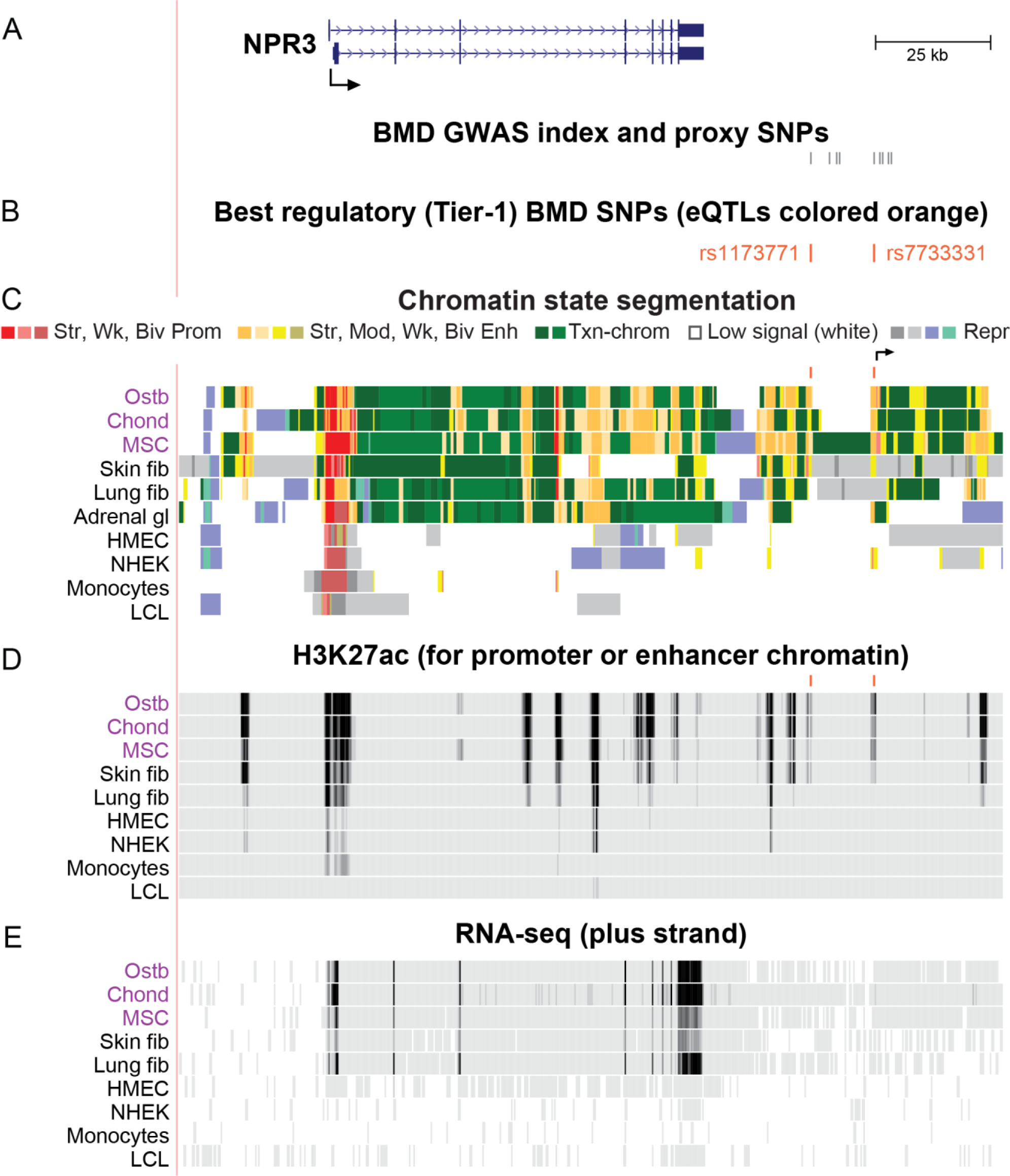
*NPR3*, which codes for a natriuretic peptide clearance receptor, is associated with two downstream Tier-1 SNPs. *(A) NPR3* (chr5:32,678,200-32,856,800) and the BMD GWAS index and proxy SNPs (*r*^2^ ≥ 0.8, EUR). *(B)* Tier-1 SNPs. *(C)* and *(D)* Chromatin state segmentation and H3K27ac profiles as in Fig. 2. *(E)* Strand-specific RNA-seq (scale, 0 – 300). Broken arrow in *(C)*, ostb/chond/MSC-specific TSS deduced from the CAGE signal for TSS; see Supplemental Fig. S8). Adrenal gl, adrenal gland; NHEK, skin keratinocytes.

The *NPR3* Tier-1 SNPs (rs1173771 and rs7733331) were downstream of the gene, in high LD (*r*^2^ = 0.99), and 14 kb apart (Fig. 3B). Both overlapped nucleosome-depleted subregions of enhancer or mixed enhancer/promoter chromatin seen preferentially in ostb, chond, and MSC (Supplemental Fig. S8D - F). An ostb-, MSC-, and chond-specific TSS for a gene encoding an unannotated long intergenic noncoding RNA (lincRNA) was found only 0.3 kb from rs7733331 by 5’ cap analysis gene expression (CAGE; Fig. 3C, arrow; Supplemental Fig. S8B). This moderate-to-weak RNA-seq signal from the plus-strand was not seen when low-sensitivity settings were used (Fig. 3E) but was seen at higher sensitivity settings specifically in ostb, MSC, and especially chond (Supplemental Fig. S8C). These SNPs were eQTLs for *NPR3* (*p* = 4 × 10^−41^ and 6 × 10^−39^) with positive effect sizes in tibial nerve, which moderately expresses this gene (Supplemental Fig. S4B). The trait-decreasing alleles of rs1173771 and rs7733331, and not the opposite alleles, are predicted to bind SATB1 (like *BICC1*’s Tier-1 SNP rs1896245) and GTF2I, respectively (Table 1). Although GTF2I is characterized as a general TF, it binds specifically to initiator and E-box DNA sequence elements in promoters^(30)^ and is implicated in osteoblastogenesis from studies on ostb and mice with heterozygous deletion of the corresponding gene (Supplemental Table S6). Therefore, rs7733331 is an especially attractive candidate for a BMD-regulatory SNP that might affect enhancer-associated *NPR3* transcription through a novel lincRNA gene in *cis*.^(34)^

### *LGR4* is associated with a BMD-risk Tier-1 SNP downstream of another protein-coding gene

Leucine rich G protein-coupled receptor 4 (*LGR4/GPR48*) was associated with one Tier-1 SNP, rs10835153. This gene encodes a cell membrane receptor that can regulate Wnt signaling and is implicated in both embryonic bone development and postnatal bone remodeling.^(35)^ Rs10835153 and most of the other BMD-associated SNPs in this region were far downstream of *LGR4* and closer to the little-studied *CCDC34* protein-encoding gene and to *BBOX1*-*AS1* than to the *LGR4* TSS (Fig. 4C) but neither of these is appreciably expressed in ostb, MSC, or chond (Supplemental Table S7). Chromatin interaction (Hi-C) profiles^(36)^ for *LGR4*-expressing lung fib and HMEC indicate that the *LGR4* promoter and the ostb enhancer chromatin overlapping *LGR4*’s intergenic Tier-1 SNP, which are 180 kb apart, are in the same large chromatin loop (topologically associated domain, TAD) ending 2 - 13 kb upstream of the *LGR4* TSS in these cells (Fig. 4B). Ostb-specific enhancer chromatin at the Tier-1 SNP and ostb/MSC/chond/HMEC/fib enhancer chromatin further downstream of *LGR4* in this TAD are likely to up-regulate the *LGR4* promoter because of the similarity in the cell type-specific enhancer chromatin profiles and *LGR4* transcription profiles (Fig. 4D and E). The trait-decreasing Alt allele of this Tier-1 SNP is predicted to bind specifically to BPTF, the largest subunit of NURF, a nucleosome remodeling factor.^(30)^ Abnormal phenotypes associated with BPTF in humans and mice include short stature and skeletal abnormalities (Supplemental Table S6).

**Figure 4.**
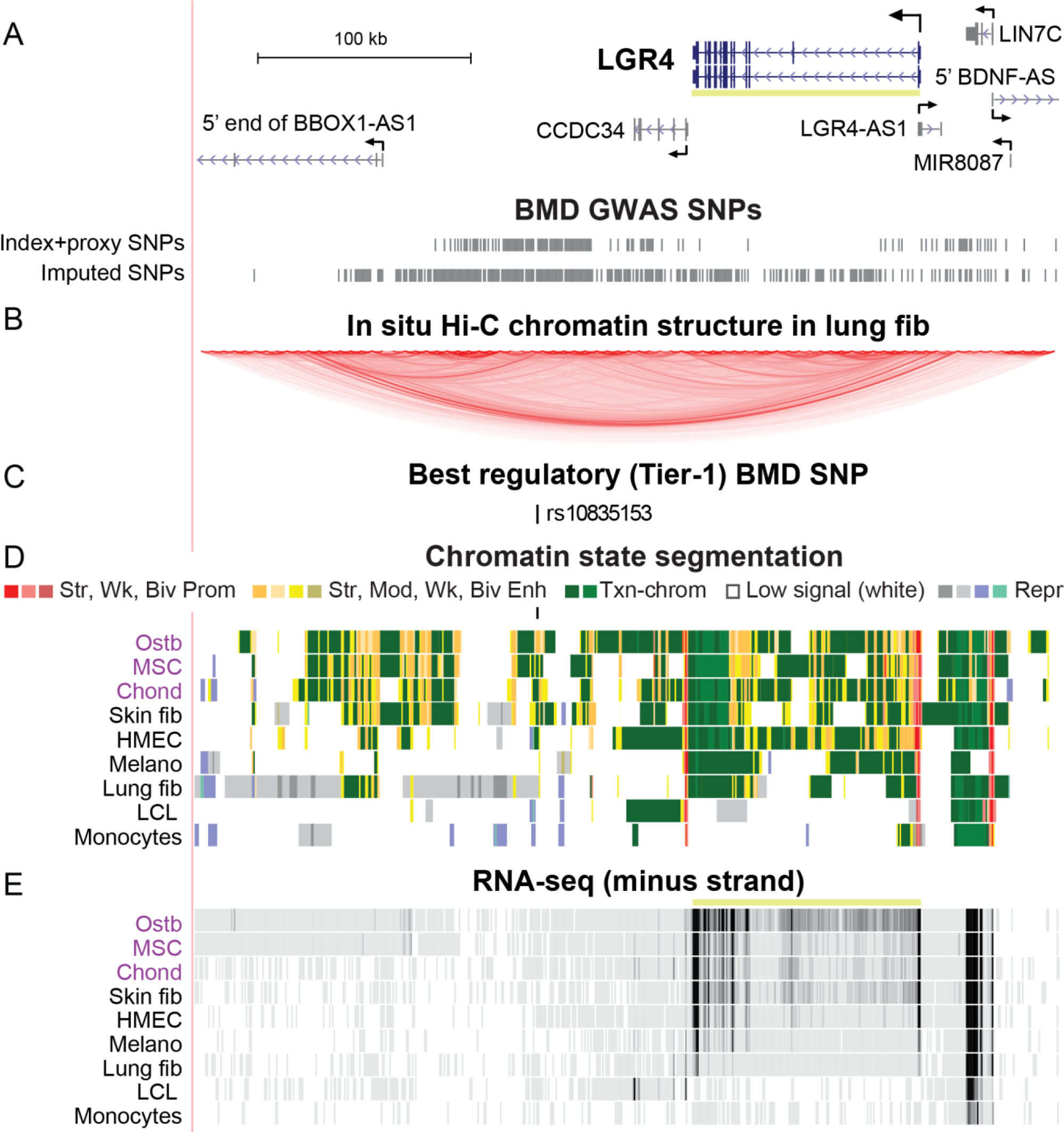
*LGR4*, which encodes a Wnt signaling receptor, has an intergenic Tier-1 SNP far downstream of the gene. *(A) LGR4* and surrounding genes (chr11:27,154,000-27,560,000) and the BMD GWAS index and proxy SNPs as well as imputed SNPs. *(B)* The Hi-C track^(36)^ for chromatin folding for the fetal lung fib cell line IMR-90; HMEC gave a similar profile (not shown). *(C)* – *(E)* as in Fig. 3 except that the minus-strand signal is shown for RNA-seq (scale, 0 – 100). Melano, foreskin melanocytes.

### *HMGA2* has three far-upstream BMD-risk Tier-1 SNPs near a novel lincRNA gene

We identified *HMGA2*, which encodes high mobility group AT-hook 2 protein, as a prime osteoporosis-risk candidate by GO term enrichment for skeletal system genes rather than by preferential expression in ostb (Fig. 1). HMGA2 is a component of multisubunit enhancer complexes (enhanceosomes). Despite *HMGA2*’s relationship to bone development^(37, 38)^ and its ostb-associated super-enhancer (a strong, unusually long enhancer) spanning the gene (Fig. 5B, dotted pink line), the RPKM ratio for ostb vs. the median of 11 non-ostb cell cultures other than MSC and chond was only of only 0.3, and the RPKM for ostb was only 0.5. Even some BMD GWAS-associated genes with negligible expression in the studied human ostb sample were linked to a limb phenotype in mouse models, possibly through effects on osteoclasts, MSC, chond, or on ostb or pre-ostb *in vivo* but not *in vitro* (Supplemental Table S8). Similarly, although the steady-state *HMGA2* RNA levels were low in the studied ostb, they were high in MSC, which can differentiate into ostb (Fig. 5C; Supplemental Table S7).

**Figure 5.**
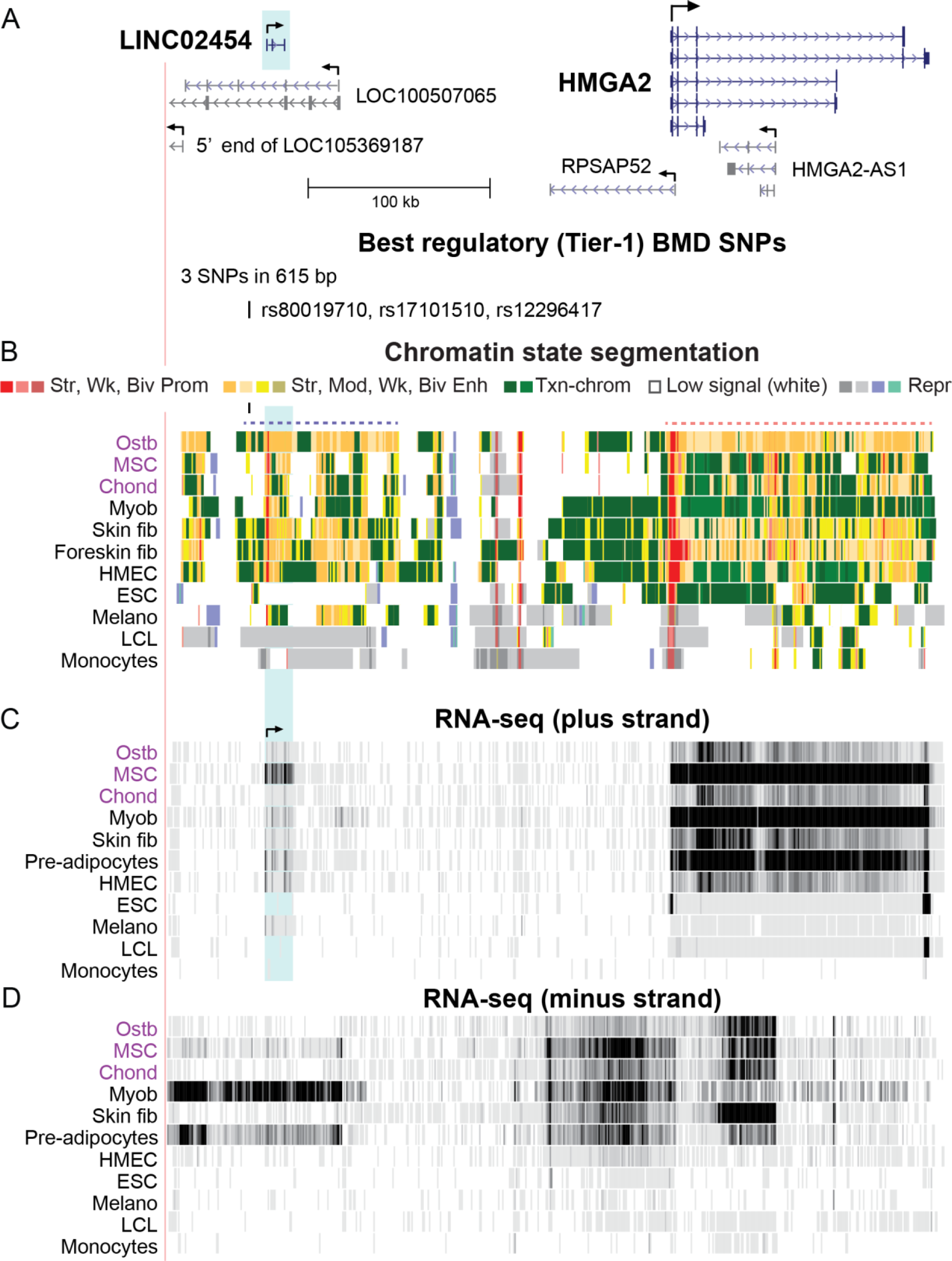
*HMGA2*, which encodes an enhancesome protein, is associated with a cluster of far-upstream intergenic Tier-1 SNPs. *(A) HMGA2* and its upstream lincRNA genes and associated Tier-1 SNPs are shown (chr12:65,942,326-66,365,457). *(B)* - *(D)* as in previous figures except that the RNA signal from both the plus- and minus-strands is shown (scales, 0 - 90 and 0 - 30, respectively). Dotted lines in panel C indicate super-enhancers in ostb; blue highlighting, *LINC02454* region. Myob, myoblasts.

*HMGA2* was associated with three intergenic Tier-1 SNPs (rs80019710, rs17101510, and rs12296417; Fig. 5A) in a 0.6-kb cluster 200 kb upstream of the *HMGA2* TSS. They were embedded in a ∼85-kb intergenic super-enhancer in ostb and a skin fib cell strain (Fig. 5B, dotted purple line), both of which had low, but appreciable, levels of *HMGA2* RNA (Fig. 5C). Within this intergenic super-enhancer are several lincRNA genes, one of which, *LINC02454*/*RP11-221N13.3*, is 9 kb downstream of the clustered Tier-1 SNPs. Its TSS overlaps a constitutive binding site for the looping protein CCCTC-binding factor (CTCF) and is surrounding by ostb/myob/skin fib-associated CTCF sites (Supplemental Fig. S9) that were not seen in cell cultures in which *HMGA2* was repressed. This suggests that the subregion containing the Tier-1 SNP cluster is involved in expression-related chromatin looping. Expression patterns of *LINC02454* and several other non-coding RNA (ncRNA) genes in this gene neighborhood were correlated with those of *HMGA2* (Fig. 5C and D and Supplemental Fig. S10). One of these ncRNA genes, *RPSAP52*, can regulate *HMGA2* both post-transcriptionally by blocking *HMGA2*-targeted miRNA activity and transcriptionally.^(39)^

From our TFBS analysis, we predict that the three *HMGA2* far-upstream Tier-1 SNPs are in DNA sequences that bind HMGA1, YY1, SATB1 or ZBTB20 specifically at their Alt alleles (Table 1). These TFs are associated with bone or cartilage biology (Supplemental Table S6). Of special interest is HMGA1, a chromatin architectural protein similar to HMGA2, which is post-transcriptionally regulated by some of the same miRNAs that control *HMGA2* RNA^(40)^ and can up-regulate Wnt signaling.^(41)^ Moreover, HMGA1 is critical for forming and maintaining bone and downregulates *miR-196A-2*, which targets *HMGA2* RNA in mouse embryonic fib.^(42)^

### Slightly varying search parameters for BMD-risk SNPs revealed likely osteoporosis-risk regulatory SNPs for *DAAM2*

By slightly relaxing our standard protocol for identifying Tier-1 SNPs, we identified three clustered SNPs (rs2504105, rs2504104, and rs2504103) associated with *DAAM2* (Dishevelled associated activator of morphogenesis 2) as highly credible candidates for BMD-associated regulatory SNPs (Fig. 6). In this modified protocol, we allowed BMD GWAS-derived EnhPro SNPs to overlap a H3K27ac narrow peak in ostb and in no more than four (rather than the previous three) of the 12 other examined cell cultures from the Roadmap database but retained all the other requirements for Tier-1 SNPs (Fig. 1). *DAAM2* encodes a key regulator of the Wnt signaling pathway and is implicated in ostb mineralization as well as in bone resorption by murine osteoclasts.^(6, 43)^ It has a ratio of 21 for ostb/non-ostb RNA levels (Supplementary Table S2) although its expression in chond and skin fib is higher than in ostb (Fig. 6E).

**Figure 6.**
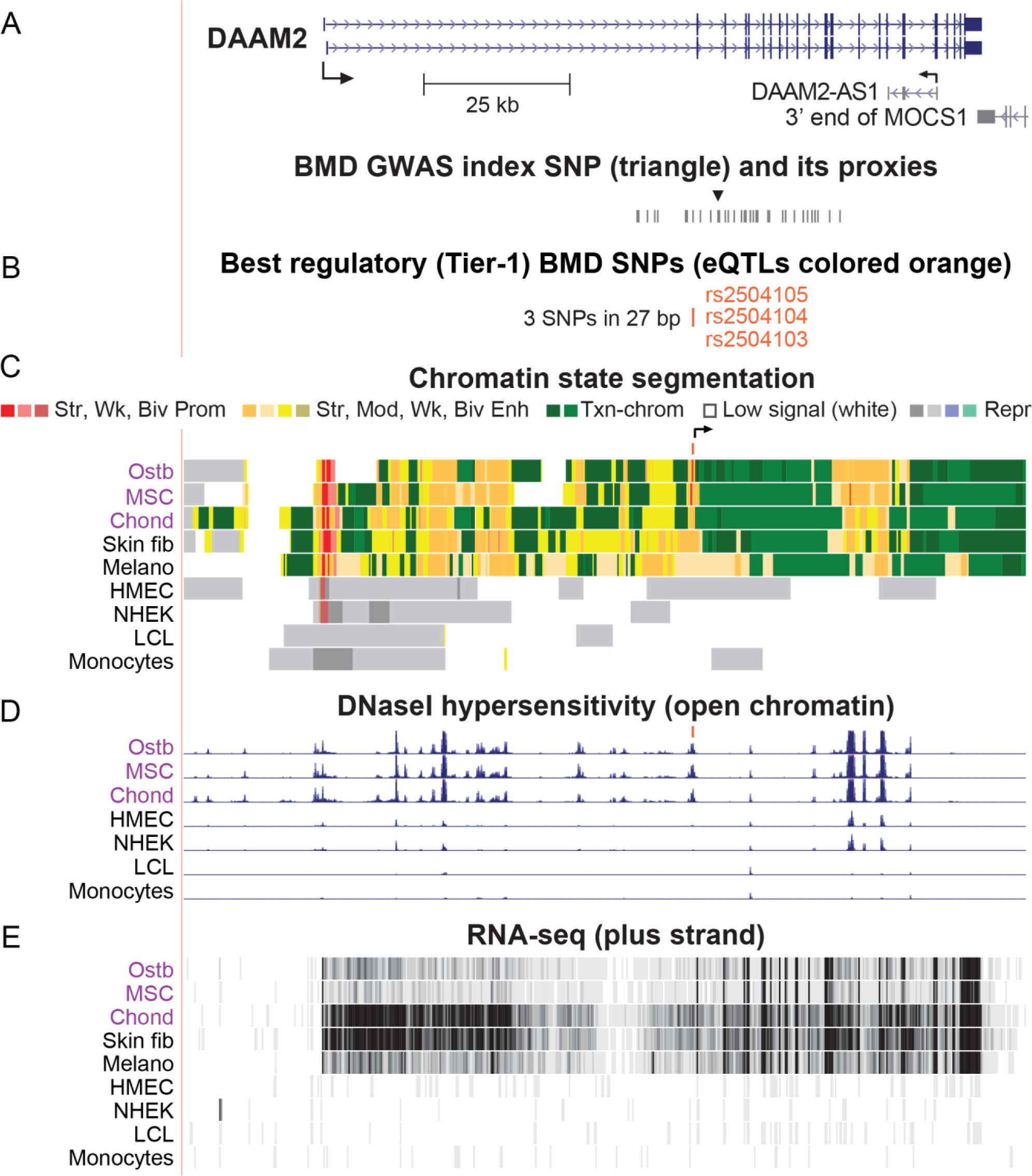
*DAAM2*, which codes for a Wnt signaling protein, has a cluster of three intragenic Tier-1 SNPs. *(A) DAAM2* and its index and proxy SNPs from BMD GWAS (chr6:39,735,841-39,911,256). *(B)* – *(E)* as in previous figures except that the scale for RNA-seq was 0 – 50; NHEK, skin keratinocytes.

Only one eBMD index SNP (rs2504101) was reported for *DAAM2* in a study by Morris et al.,^(6)^ who featured this gene as a candidate causal gene for osteoporosis risk. We found that their selected index SNP in intron 2 of *DAAM2* (Fig. 6A, triangle) overlaps transcription-type (H3K36me3-enriched) chromatin and not enhancer or promoter chromatin in ostb, MSC, chond, and skin fib and does not overlap a DHS in any examined cell type. From 38 SNPs in high LD with this index SNP (*r*^2^ > 0.8, EUR; Supplemental Table S3), we identified three (rs2504105, rs2504104, and rs2504103; LD *r*^2^ = 0.97) as Tier-1 SNPs using the above-described criterion for H3K27ac specificity. These SNPs span only 27 bp and were embedded in mixed enhancer/promoter chromatin in ostb and enhancer chromatin or enhancer/promoter chromatin in MSC, chond, and skin fib. They were located 0.8 kb upstream of exon 2 of *DAAM2* and 0.4 kb upstream of a novel sense-strand intronic TSS (CAGE signal; Fig. 6C, arrow) in these cell types but not in cell cultures with repressed *HMGA2*. Moreover, they are eQTLs for tibial nerve (*p* = 8.4 × 10^−6^), a *DAAM2-*expressing tissue (Supplemental Fig. S4D). These findings suggest that the cluster of Tier-1 SNPs up-regulates *DAAM2* expression partly by upregulating expression of an intronic ncRNA transcript at a cell type-specific enhancer. The much lower steady-state levels of *DAAM2* RNA in ostb and MSC than in chond despite their similar amounts of enhancer chromatin could be due to post-transcriptional regulation.

Like rs11006188 in *BICC1*, the Ref allele of *DAAM2* Tier-1 SNP rs2504105 is predicted to bind SMAD family TFs (Supplemental Table S6). In addition, the Alt allele of Tier-1 SNP rs2504104 overlaps predicted binding sites for RUNX2, the master regulator of ostb lineage commitment,^(11)^ and CBFB, which can help stabilize Runx-family proteins binding to DNA^(44)^ (Table 1). Lastly, the Alt allele of rs2504103 is predicted to bind NFIC, which plays important roles in both tooth and bone development.^(45)^ These findings support the candidacy of one or more of these SNPs as causal regulatory SNPs for BMD risk.

## Discussion

We identified 16 high-priority osteoporosis-risk regulatory SNPs by using an unusual in-depth bioinformatic approach relying on epigenomic and transcriptomic comparisons of ostb and heterologous cell cultures as well as prediction of allele-specific binding of TFs that are important for bone biology. These Tier-1 SNPs, which had not been previously identified as candidates for causal regulatory SNPs for BMD, were associated with five skeletal system-related genes. Three of these, *BICC1, LGR4, DAAM2*, are involved directly or indirectly in canonical Wnt signaling, a signaling pathway that plays critical roles in osteoblastogenesis and bone homeostasis.^(46-48)^ The other two genes are *NPR3*, and *HMGA2*, a mediator of natriuretic signaling and an essential component of enhanceosomes, respectively.^(28, 49^)

One of the most interesting genes for which we identified novel regulatory candidate SNPs is *BICC1*, which encodes a self-polymerizing RNA-binding protein that indirectly regulates Wnt signaling.^(32)^ This gene had two clusters of Tier-1 SNPs near its 5’ end (Fig. 2). BICC1 is involved in osteoblastogenesis and polycystic kidney disease, in part, through its inhibition of post-transcriptional silencing of *PKD2* RNA by *miR-17*.^(31, 50, 51)^ PKD2/PC2 and PKD1/PC1 form primary cilia that function as osteoporosis-related mechanosensors in ostb (and probably osteocytes) and in epithelial kidney cells, and BICC1 has been detected in primary cilia in kidney cells.^(32, 52-54)^ Conditional inactivation of *Pkd2* in ostb/osteocytes of mice results in decreased levels of *Runx2* RNA and Wnt signaling along with the development of osteopenia.^(51)^ An important study by Mesner et al.^(31)^ implicating BICC1 in osteoporosis demonstrated that a heterozygous inactivating mutation in *Bicc1* in male mice led to low femoral BMD and that *Bicc1*-deficent ostb had impaired differentiation *in vitro* that could be rescued by *Pkd2* overexpression.

*LGR4*, which had one associated Tier-1 SNP (Fig. 4), encodes a bone-related Wnt signaling receptor^(29, 55)^ and had been catalogued as one of many genes with an eBMD-associated missense SNPs (rs34804482) predicted to have a deleterious effect on protein structure.^(6)^ Moreover, Styrkarsdottir et al.^(56)^ found a rare nonsense mutation in the gene that is associated with low BMD, osteoporotic fractures in elderly individuals, electrolyte imbalance, and several types of cancer. In mice, homozygous deletion of *Lgr4* results in a low-BMD phenotype, a strong delay in ostb differentiation and mineralization, and elevated numbers of osteoclasts.^(29)^ Osteoclasts and MSC precursors of ostb have also been implicated in the positive effects of LGR4 on bone formation and homeostasis from conditional mouse knock-out models and *in vitro* studies.^(57, 58)^ However, the single Tier-1 SNP that we found to be associated with *LGR4* was in enhancer chromatin in ostb but not in MSC. Epigenetic profiles of osteoclasts are not available but the finding of negligible levels of expression of *Lgr4* (and the mouse homologs of the other Tier-1 associated genes) in cultured mouse osteoclasts and substantial levels in mouse ostb (Supplemental Fig. S6) suggests that the *LGR4* Tier-1 SNP’s regulatory role in osteoporosis risk is through ostb and possibly osteocytes.

Like *LGR4, DAAM2* encodes a regulator of canonical Wnt signaling implicated in the ostb and osteoclast lineages.^(6, 43, 59)^ Its 27-bp cluster of three intragenic Tier-1 SNPs, which are in perfect LD, may act through their overlapping enhancer/promoter chromatin in ostb, MSC, and chond and the novel intragenic TSS immediately downstream of these SNPs (Fig. 6C, arrow). Consistent with a role for *DAAM2* in inherited osteoporosis risk, an eBMD GWAS-derived missense variant (rs201229319) in *DAAM2* that is probably deleterious had been identified previously by Morris et al.^(6)^ They also found that homozygous knockout of *Daam2* in mice decreases bone strength and increases cortical bone porosity and that CRISPR/Cas9-mediated inactivation of *DAAM2* impairs mineralization in an ostb cell line. We identified the *DAAM2* Tier-1 SNPs by a very small change in one of our requirements for Tier-1 SNP designation (Fig. 1) to < 5 rather than < 4 of the 12 non-ostb related cell cultures displaying the ostb-associated H3K27ac peak overlapping the SNP. Therefore, future bioinformatic studies could uncover many more plausible BMD-regulatory SNPs by small variations of our protocol.

*NPR3 (NPRC)*, one of the Tier-1 SNP associated genes, is involved not only in linear bone growth, bone turnover, and endochondral ossification,^(28, 60-62)^ but also in cardiovascular homeostasis, and renal cancer metastasis.^(28, 62)^ Its Tier-1 SNPs (rs1173771 and rs7733331), which are in almost perfect LD, are upstream of another novel TSS observed preferentially in ostb and chond (Fig. 3C and Supplemental Fig. S8). *NPR3* encodes a transmembrane clearance receptor that opposes natriuretic peptide signaling through NPR1 and NPR2 receptors, which, unlike NPR3, contain guanylyl cyclase activity that is activated upon peptide binding. Enhancement of bone growth was found in patients with bi-allelic *NPR3* loss-of-function mutations or monoallelic gain-of-function mutations in *NPR2*.^(28)^ Osteocrin, a protein structurally resembling natriuretic peptides, binds specifically to NPR3 and is implicated in stimulating bone growth by limiting NPR3-mediated clearance of natriuretic peptides.^(63)^ *NPR3* was associated with high bone density in a GWAS.^(64)^ *NPR3* is downregulated post-transcriptionally by *miR-143*,^(65)^ whose host gene, *CARMN*, displays especially high expression in ostb.^(66)^ Both *NPR3*-associated Tier-1 SNPs have been previously identified as index SNPs in many blood pressure-related GWAS, consistent with the role of *NPR3* also in the cardiovascular system.^(19)^ In addition, rs1173771 was associated with height in a GWAS.^(67)^

*HMGA2*, which has three clustered Tier-1 SNPs far upstream of the gene (Fig. 5), was the only one of the five prioritized genes that was identified by its bone-related GO association rather than by its ostb/non-ostb expression ratio (Fig. 1). Some genes that impact osteoporosis risk through the ostb lineage may act at the pre-ostb or osteocyte stages. Alternatively, some osteoporosis-associated genes might be negatively associated with osteoblastogenesis. *HMGA2* may be an example of both phenomena. It encodes a chromosomal architectural and enhanceosomal protein implicated in negatively regulating the differentiation of bone marrow-derived MSC to ostb^(68)^ and is necessary for normal embryogenesis, including bone development.^(37, 38)^ It also is involved in carcinogenesis and metastasis,^(69)^ cell replication, and autophagy, which plays central roles in bone homeostasis through effects on ostb, osteocytes, and osteoclasts.^(70)^ *HMGA2* displays very low or negligible expression in normal postnatal tissues but is highly expressed in some mesoderm-derived stem or progenitor cell types including MSC, myob, and pre-adipocytes. It is much more weakly expressed in ostb, in accord with its negative effects on ostb formation from MSC.^(49)^ The need for tight control of *HMGA2* expression is evidenced by its unusually complex regulatory circuitry including the cancer-associated *let-7* miRNA and the *HMGA2*-overlapping pseudogene *RPSAP52* and anti-sense gene *HMGA2-AS1.*^(49, 71)^ Importantly, another ncRNA gene, *LINC02454*, which exhibits an *HMGA2-*like expression pattern (Supplemental Fig. S10), is 9 kb downstream of the three Tier-1 SNPs in an ostb-associated super-enhancer far from *HMGA2. LINC02454* was previously referenced only for its dysregulation in cancer cells.^(72)^ We propose that one or several of these Tier-1 SNPs help control expression of *LINC02454*, which, in turn, helps regulate transcription of *HMGA2* in MSC. The much lower steady-state levels of *HMGA2* mRNA in ostb than in MSC despite the abundant enhancer chromatin within or upstream of the gene in ostb (Fig. 5) may be due to multiple miRNAs targeting *HMGA2*.^(71)^ Besides a role for *HMGA2* in negatively regulating MSC differentiation to ostb, it may play a protective role in the skeleton^(73)^ at later steps in differentiation of ostb to osteocytes and in osteocyte homeostasis by helping to induce autophagy.^(70, 74, 75)^

Like our five osteoporosis-risk candidate genes, the TFs predicted to bind to the Tier-1 SNP-containing DNA sequences associated with these genes have special functions in the skeletal system (Supplemental Table S6). Further evidence implicating Tier-1 SNPs as transcription-regulatory variants comes from our finding that 11 of the 16 SNPs for *BICC1, DAAM2*, or *NPR3* overlapped eQTLs for one or several tissues in which these genes are expressed. However, eQTLs are not available for ostb and, given our requirement that Tier-1 SNPs be present in enhancer or promoter chromatin preferentially in ostb, eQTL overlap of Tier-1 SNPs should be underrepresented by available data. In the cases of *NPR3*, and *DAAM2*, which had 2 - 3 Tier-1 SNPs in high LD, only a single SNP at each of these loci may be the regulatory causal SNP. However, for *BICC1*, the dichotomy of the direction of effect sizes for tissue eQTLs and the distribution of BMD-increasing alleles for the two clusters of Tier-1 SNPs suggests that at least one Tier-1 SNP in the TSS-adjacent cluster and one in the 60-kb downstream cluster are causal regulatory SNPs.

## Conclusion and future directions

Candidates for highly credible causal regulatory GWAS-derived variants are more difficult to prioritize but are much more numerous than exonic variants affecting polypeptide structure. For our selection of 16 osteoporosis-related GWAS regulatory SNPs associated with five genes from 38 BMD GWAS, we used very restrictive epigenetic and transcriptomic criteria. The approach we took to prioritize these osteoporosis-associated genes and their Tier-1 SNPs for future experimental studies can be extended for additional prioritization of candidates for regulatory BMD SNPs, for example, by changing the threshold for preferential expression in ostb, using different tools for gene functional analysis, making small changes in the requirements for designation of preferential enhancer or promoter chromatin in ostb, looking for preferential expression in chond instead of ostb, and expanding the analyses of epigenomic and transcriptomic profiles to osteoclasts and osteocytes as these profiles become available. Elucidation of regulatory genetic variants strongly associated with disease-susceptibility using epigenetic-intensive strategies like ours can help find marker SNPs to identify at-risk individuals for treatment or life-style modifications and to develop pharmacological interventions by focusing attention on insufficiently studied protein-coding genes and neighboring ncRNA genes.

## Disclosures

All of the authors state that they have no conflict of interest.

## Acknowledgements

This study was partially supported or benefited by grants from the National Institutes of Health [P20GM109036, R01AR069055, U19AG055373, and R01MH104680], and the Edward G. Schlieder Endowment and Drs. W. C. Tsai and P. T. Kung Professorship in Biostatistics from Tulane University.

## Authors contributions

XZ and ME designed the study. H-WD and HS gave advice on the study design. XZ performed analysis of the data and prepared figures and tables. ME and XZ wrote the manuscript. All authors revised and reviewed the paper.

## Notes

### Competing Interest Statement

The authors have declared no competing interest.

