## Supplemental Table S1-S8 for "Prioritization of osteoporosis-associated GWAS SNPs using epigenomics and transcriptomics"

**Supplemental Table S1. The 38 BMD GWA studies used for this analysis[a]**

| First author | Study | BMD region/type[b] | Ancestry population | Year | PMID |
|---|---|---|---|---|---|
| Chesi A | A trans-ethnic genome-wide association study identifies gender-specific loci influencing pediatric aBMD and BMC at the distal radius. | R | EUR & non-EUR | 2015 | 26041818 |
| " | A genomewide association studyidentifies two sex-specific loci, at SPTB and IZUMO3, influencing pediatric bone mineral density at multiple skeletal sites. | FN, H, R, S | EUR only | 2017 | 28181694 |
| Choi HJ | Genome-wide association study in East Asians suggests UHMK1 as a novel bone mineral density susceptibility gene. | FN, H, LS | EUR & non-EUR | 2016 | 27424934 |
| Duncan EL | Genome-wide association study using extreme truncate selection identifies novel genes affecting bone mineral density and fracture risk. | H | EUR only | 2011 | 21533022 |
| Estrada K | Genome-wide meta-analysis identifies 56 bone mineral density loci and reveals 14 loci associated with risk of fracture. | FN, LS | EUR & non-EUR | 2012 | 22504420 |
| Gregson CL | Genome-wide association study of extreme high bone mass: Contribution of common genetic variation to extreme BMD phenotypes and potential novel BMD-associated genes. | H, LS | EUR & non-EUR | 2018 | 29883787 |
| Kemp JP | Phenotypic dissection of bone mineral density reveals skeletal site specificity and facilitates the identification of novel loci in the genetic regulation of bone mass attainment. | Limb, skull, TBLH | EUR & non-EUR | 2014 | 24945404 |
| " | Identification of 153 new loci associated with heel bone mineral density and functional involvement of GPC6 in osteoporosis. | Heel | EUR only | 2017 | 28869591 |
| Kichaev G | Leveraging polygenic functional enrichment to improve GWAS power. | Heel | EUR only | 2018 | 30595370 |
| Kim SK | Identification of 613 new loci associated with heel bone mineral density and a polygenic risk score for bone mineral density, osteoporosis and fracture. | Heel | EUR only | 2018 | 30048462 |
| Kung AW | Association of JAG1 with bone mineral density and osteoporotic fractures: a genome-wide association study and follow-up replication studies. | FN, LS | EUR & non-EUR | 2010 | 20096396 |
| Liang X | Assessing the genetic correlations between early growth parameters and bone mineral density: a polygenic risk score analysis. | FN, forearm, H, S, TB | EUR only | 2018 | 30172743 |
| Lu S | Bivariate genome-wide association analyses identified genetic pleiotropic effects for bone mineral density and alcohol drinking in Caucasians. | H, S, TB | EUR only | 2016 | 28012008 |
| Medina-Gomez C | Meta-analysis of genome-wide scans for total body BMD in children and adults reveals allelic heterogeneity and age-specific effects at the WNT16 locus. | Skull, TBLH | EUR only | 2012 | 22792070 |
| " | Bivariate genome-wide association meta-analysis of pediatric musculoskeletal traits reveals pleiotropic effects at the SREBF1/TOM1L2 locus. | TBLH | EUR only | 2017 | 28743860 |
| " | Life-course genome-wide association study meta-analysis of total body BMD and assessment of age-specific effects. | TBLH | EUR & non-EUR | 2018 | 29304378 |
| Mitchell JA | Multi-dimensional bone density phenotyping reveals new insights in to genetic regulation of the pediatric skeleton. | FN, H, LS, R | EUR & non-EUR | 2017 | 29240982 |
| Moayyeri A | Genetic determinants of heel bone properties: genome-wide association meta-analysis and replication in the GEFOS/GENOMOS consortium. | Heel | EUR only | 2014 | 24430505 |
| Morris JA | An atlas of genetic influences on osteoporosis in humans and mice. | Heel | EUR only | 2018 | 30598549 |
| Mullin BH | Genome-wide association study using family-based cohorts identifies the WLS and CCDC170/ESR1 loci as associated with bone mineral density. | FN, H, LS | EUR only | 2016 | 26911590 |
| Nielson CM | Novel genetic variants associated with increased vertebral volumetric BMD, reduced vertebral fracture risk, and increased expression of SLC1A3 and EPHB2. | LS (vBMD) | EUR only | 2016 | 27476799 |
| Paternoster L | Genome-wide association meta-analysis of cortical bone mineral density unravels allelic heterogeneity at the RANKL locus and potential pleiotropic effects on bone. | FN, LS, TB | EUR only | 2010 | 21124946 |
| " | Genetic determinants of trabecular and cortical volumetric bone mineral densities and bone microstructure. | vBMD | EUR only | 2013 | 23437003 |
| Pei YF | Association of 3q13.32 variants with hip trochanter and intertrochanter bone mineral density identified by a genome-wide association study. | H | EUR & non-EUR | 2016 | 27311723 |
| " | Genome-wide association meta-analyses identified 1q43 and 2q32.2 for hip Ward's triangle areal bone mineral density. | FN | EUR & non-EUR | 2016 | 27397699 |
| " | Joint study of two genome-wide association meta-analyses identified 20p12.1 and 20q13.33 for bone mineral density. | FN, LS | EUR & non-EUR | 2018 | 29499414 |
| Richards JB | Bone mineral density, osteoporosis, and osteoporotic fractures: a genome-wide association study. | FN, LS | EUR only | 2008 | 18455228 |
| Rivadeneira F | Twenty bone-mineral-density loci identified by large-scale meta-analysis of genome-wide association studies. | FN, LS | EUR only | 2009 | 19801982 |
| Styrkarsdottir U | Multiple genetic loci for bone mineral density and fractures. | FN, H, LS | EUR only | 2008 | 18445777 |
| " | New sequence variants associated with bone mineral density. | H, S | EUR only | 2008 | 19079262 |
| " | Nonsense mutation in the LGR4 gene is associated with several human diseases and other traits. | H, LS, TB | EUR only | 2013 | 23644456 |
| " | Sequence variants in the PTCH1 gene associate with spine bone mineral density and osteoporotic fractures. | H, LS | EUR & non-EUR | 2016 | 26733130 |
| Tan LJ | Bivariate genome-wide association study implicates ATP6V1G1 as a novel pleiotropic locus underlying osteoporosis and age at menarche. | H, LS | EUR & non-EUR | 2015 | 26312577 |
| Xiong DH | Genome-wide association and follow-up replication studies identified ADAMTS18 and TGFBR3 as bone mass candidate genes in different ethnic groups. | H, S | EUR only | 2009 | 19249006 |
| Zhang L | Multistage genome-wide association meta-analyses identified two new loci for bone mineral density. | FN, H, LS | EUR & non-EUR | 2013 | 24249740 |
| Zheng HF | WNT16 influences bone mineral density, cortical bone thickness, bone strength, and osteoporotic fracture risk. | Forearm | EUR only | 2012 | 22792071 |
| " | Meta-analysis of genome-wide studies identifies MEF2C SNPs associated with bone mineral density at forearm. | Forearm | EUR & non-EUR | 2013 | 23572186 |
| " | Whole-genome sequencing identifies EN1 as a determinant of bone density and fracture. | FN, forearm, S | EUR only | 2015 | 26367794 |

[a]All studies were from the GWAS catalogue: https://www.ebi.ac.uk/gwas/

[b]FN, femoral neck; H, total hip; LS, lumbar spine; R, radius; S, total spine; TB, total body; TBLS, total body less head; vBMD, volumetric BMD (g/cm$^3$)

**Supplemental Table S2. Expression ratios for some reference genes associated with BMD GWAS SNPs from the studies in Table S1: RNA-seq data for expression in osteoblasts, mesenchymal stem cells, or chondrocytes vs. in heterologous cell cultures[a]**

| BMD GWAS reference genes | Ratio of RPKM in ostb, MSC or chond to the median RPKM of 11 heterologous cell cultures[c] | | | Average RPKM from technical duplicates (ENCODE database) | | | BMD GWAS reference genes | Ratio of RPKM in ostb, MSC or chond to the median RPKM of 11 heterologous cell cultures[c] | | | Average RPKM from technical duplicates (ENCODE database) | | |
|---|---|---|---|---|---|---|---|---|---|---|---|---|---|
| | Ostb/non-ostb (excl. MSC & chond) | MSC/non-MSC (excl. ostb & chond) | Chond/non-chond (excl. ostb & MSC) | Ostb | MSC | Chond | | Ostb/non-ostb (excl. MSC & chond) | MSC/non-MSC (excl. ostb & chond) | Chond/non-chond (excl. ostb & MSC) | Ostb | MSC | Chond |
| **DAAM2**[b] | **20.61** | **6.82** | **34.59** | **1.23** | **0.41** | **2.07** | SATB2 | 4.53 | 8.99 | 0.30 | 2.28 | 4.53 | 0.15 |
| **NPR3** | **14.61** | **7.21** | **22.31** | **3.17** | **1.57** | **4.85** | ZFHX4 | 6.28 | 6.65 | 2.37 | 2.27 | 2.40 | 0.86 |
| **BICC1** | **5.26** | **4.68** | **5.26** | **7.14** | **6.34** | **7.13** | GPC6 | 9.35 | 6.75 | 22.32 | 2.22 | 1.60 | 5.30 |
| **LGR4** | **5.13** | **3.38** | **2.19** | **3.03** | **2.00** | **1.29** | CSF1 | 1.66 | 6.41 | 2.76 | 2.13 | 8.22 | 3.53 |
| **HMGA2** | **0.30** | **2.43** | **0.25** | **0.53** | **4.20** | **0.44** | METTL7A | 2.18 | 5.90 | 3.48 | 2.13 | 5.76 | 3.40 |
| TBX15 | 296.80 | 410.35 | 479.88 | 2.07 | 2.86 | 3.34 | EBF1 | 18.66 | 12.12 | 25.81 | 2.12 | 1.37 | 2.92 |
| ADAM12 | 16.08 | 4.57 | 8.20 | 13.29 | 3.78 | 6.78 | APOL1 | 5.07 | 4.46 | 4.32 | 1.85 | 1.63 | 1.58 |
| SPECC1 | 8.64 | 2.30 | 4.53 | 6.50 | 1.73 | 3.41 | CCBE1 | 6.62 | 7.84 | 10.04 | 1.70 | 2.01 | 2.57 |
| RUNX2 | 6.07 | 4.14 | 0.21 | 1.15 | 0.78 | 0.04 | CD68 | 1.73 | 5.30 | 4.33 | 1.61 | 4.92 | 4.03 |
| COL1A1 | 6.32 | 1.02 | 1.18 | 164.90 | 26.59 | 30.83 | NEGR1 | 1.99 | 8.38 | 6.80 | 1.51 | 6.36 | 5.16 |
| MMP2 | 8.69 | 11.77 | 1.47 | 77.28 | 104.63 | 13.03 | FZD1 | 2.85 | 6.32 | 0.86 | 1.47 | 3.26 | 0.44 |
| LRP1 | 11.57 | 9.67 | 5.76 | 65.12 | 54.43 | 32.43 | HOXA10 | 5.37 | 8.79 | 5.20 | 1.17 | 1.92 | 1.13 |
| CDH11 | 12.62 | 6.98 | 4.28 | 59.09 | 32.68 | 20.03 | SLC4A4 | 84.36 | 120.07 | 33.45 | 1.15 | 1.63 | 0.45 |
| GALNT1 | 9.22 | 3.26 | 5.78 | 31.40 | 11.11 | 19.66 | MMP16 | 2.55 | 6.53 | 0.32 | 1.08 | 2.77 | 0.14 |
| TMEM119 | 26.88 | 16.24 | 2.37 | 14.64 | 8.84 | 1.29 | AKR1C2 | 5.90 | 22.22 | 20.29 | 1.07 | 4.01 | 3.66 |
| PRRX1 | 2.92 | 5.77 | 3.91 | 12.45 | 24.65 | 16.68 | LMOD1 | 15.68 | 4.20 | 1.99 | 1.00 | 0.27 | 0.13 |
| PPAP2B | 4.06 | 7.18 | 5.73 | 11.37 | 20.09 | 16.03 | EYA4 | 19.93 | 5.95 | 0.29 | 0.88 | 0.26 | 0.01 |
| SFRP4 | 2199.36 | 452.14 | 425.09 | 10.94 | 2.25 | 2.11 | EPSTI1 | 7.35 | 8.66 | 5.40 | 0.85 | 1.01 | 0.63 |
| LOXL1 | 3.21 | 6.14 | 2.00 | 9.46 | 18.08 | 5.90 | TGFB2 | 1.94 | 4.74 | 5.72 | 0.79 | 1.93 | 2.33 |
| COL11A1 | 16.16 | 0.13 | 10.05 | 6.43 | 0.05 | 4.00 | KIAA1671 | 3.00 | 8.00 | 2.42 | 0.79 | 2.09 | 0.63 |
| NRG1 | 6.27 | 2.17 | 0.06 | 5.81 | 2.01 | 0.05 | FAM101A | 26.05 | 1.45 | 19.97 | 0.79 | 0.04 | 0.60 |
| AKR1C3 | 3.23 | 5.26 | 1.98 | 4.49 | 7.31 | 2.75 | CACNA1C | 5.36 | 2.43 | 6.51 | 0.76 | 0.34 | 0.92 |
| VASN | 6.80 | 6.92 | 5.42 | 4.40 | 4.48 | 3.51 | EYA1 | 96.41 | 136.33 | 0.62 | 0.71 | 1.00 | 0.00 |
| SLC1A3 | 3.93 | 6.20 | 0.01 | 3.50 | 5.52 | 0.01 | SPRED2 | 1.17 | 6.13 | 1.08 | 0.65 | 3.42 | 0.60 |
| PLXDC2 | 14.13 | 6.59 | 19.25 | 3.43 | 1.60 | 4.67 | COL21A1 | 21.95 | 21.36 | 2.27 | 0.64 | 0.63 | 0.07 |
| ARID5B | 2.04 | 1.94 | 7.87 | 3.34 | 3.18 | 12.88 | ITIH4 | 10.65 | 3.25 | 4.60 | 0.63 | 0.19 | 0.27 |
| ANK3 | 7.61 | 5.37 | 2.48 | 3.30 | 2.33 | 1.07 | HOXA11 | 13.44 | 15.09 | 16.43 | 0.58 | 0.65 | 0.71 |
| SMOC1 | 26.53 | 22.09 | 58.59 | 2.95 | 2.45 | 6.51 | MAFB | 3.85 | 9.84 | 10.20 | 0.58 | 1.49 | 1.54 |
| PTGIS | 17.94 | 36.57 | 10.04 | 2.92 | 5.95 | 1.63 | PCOLCE2 | 4.22 | 21.16 | 29.00 | 0.58 | 2.90 | 3.98 |
| DEPTOR | 39.22 | 1.94 | 0.87 | 2.87 | 0.14 | 0.06 | CD4 | 13.89 | 18.06 | 3.33 | 0.55 | 0.72 | 0.13 |
| MIR143HG | 45.58 | 7.05 | 4.30 | 2.66 | 0.41 | 0.25 | ZNF423 | 10.79 | 10.04 | 1.77 | 0.53 | 0.49 | 0.09 |
| TRIB2 | 2.25 | 5.05 | 1.00 | 2.49 | 5.61 | 1.11 | PPM1H | 7.14 | 6.47 | 7.96 | 0.52 | 0.47 | 0.58 |
| TNFRSF11B | 61.76 | 9.64 | 21.44 | 2.43 | 0.38 | 0.84 | ITGB2 | 110.51 | 23.78 | 71.67 | 0.51 | 0.11 | 0.33 |
| GAS1 | 8.61 | 6.22 | 4.19 | 2.40 | 1.74 | 1.17 | C4orf22 | 9.53 | 26.24 | 0.19 | 0.50 | 1.39 | 0.01 |
| SMAD7 | 5.26 | 6.91 | 2.29 | 2.40 | 3.16 | 1.04 | | | | | | | |
| FOXC1 | 7.08 | 15.50 | 13.70 | 2.30 | 5.05 | 4.46 | | | | | | | |
| UBE2E3 | 0.85 | 5.63 | 0.62 | 2.30 | 15.16 | 1.68 | | | | | | | |

[a]From the 2,157 genes reported to be associated with 3,214 unique BMD GWAS index SNPs ($p < 5 \times 10^{-8}$) were obtained from the GWAS Catalog (as of October 2019, https://www.ebi.ac.uk/gwas); this table shows the 70 genes with reads per kilobase million (RPKM) in osteoblasts (ostb) of > 0.5 and ratios of preferential expression in ostb, mesenchymal stem cells (MSC), or chondrocytes (chond) of > 5 (see columns 3 - 5); except for *HMGA2* (see footnote b). The transcription data are from ENCODE long RNA-seq (whole cell, total RNA: ENCODE/Cold Spring Harbor lab; http://genome.ucsc.edu)

[b]The five best candidate genes for association with BMD regulatory SNPs from this study (see main text) are in bold font. The seven genes that are preferentially expressed in ostb and associated with at least one epigenetically prioritized (EnhPro, see Supplemental Table S3, footnote **a**) SNP are in blue font, except for DAAM2, for which a slightly relaxed epigenetic criterion was used (see Supplemental Table S3, footnote **c**); *HMGA2* was chosen based upon GREAT enrichment analysis (http://great.stanford.edu/) instead of overexpression in ostb

[c]Average RPKM (reads per kilobase million) for technical duplicates divided by the median expression of other 11 heterologous cell cultures, namely, skin fibroblasts, fetal lung fibroblasts (IMR-90), myoblasts, human mammary epithelial cells (HMEC), foreskin or dermal melanocytes, aortic endothelial cells, follicle dermal papilla cells, pericytes, saphenous vein endothelial cells, and preadipocytes; for HMEC only a single value was available

**Supplemental Table S3. The 154 BMD GWAS SNPs that fit the epigenetic criteria to be EnhPro SNPs[a]**

| Examined SNP (Tier-1 SNP associated gene) | Overlap with strong enh or prom chromatin segments (Roadmap Auxilliary State 1, 3, 8 or 9) or other type of chromatin | | |
|---|---|---|---|
| | Ostb | MSC | Chond |
| rs1173771[b] (NPR3) | Enh | Other | Other |
| rs7733331 (NPR3) | Enh | Enh/prom | Enh |
| rs112597538 (BICC1) | Prom | Prom | Prom |
| rs1896245 (BICC1) | Enh/prom | Enh/prom | Other |
| rs1896243 (BICC1) | Enh | Enh/prom | Other |
| rs1658429 (BICC1) | Enh | Enh | Other |
| rs4948523 (BICC1) | Enh | Enh | Enh |
| rs10835153 (LGR4) | Enh | Other | Other |
| rs17101510 (HMGA2) | Enh | Other | Other |
| rs12296417 (HMGA2) | Enh | Other | Other |
| rs2504105[c] (DAAM2) | Enh/prom | Enh/prom | Enh |
| rs2504104[c] (DAAM2) | Enh/prom | Enh | Enh |
| rs2504103[c] (DAAM2) | Enh/prom | Enh | Enh |
| rs35110415 | Enh | Other | Other |
| rs78653043 | Enh | Other | Other |
| rs12078572 | Enh | Other | Enh |
| rs1891707 | Enh | Other | Enh |
| rs6680737 | Enh/prom | Other | Other |
| rs16844133 | Enh/prom | Enh | Enh |
| rs16844134 | Enh/prom | Enh | Enh |
| rs1465488 | Enh | Other | Other |
| rs6432337 | Enh | Other | Other |
| rs6751241 | Enh | Other | Other |
| rs139168054 | Enh | Enh | Other |
| rs7609256 | Enh | Enh | Enh |
| rs17045474 | Enh | Enh | Enh |
| rs10210502 | Enh | Other | Other |
| rs6732131 | Enh | Enh | Enh |
| rs5834956 | Enh | Enh | Enh |
| rs6430092 | Enh | Other | Other |
| rs10635110 | Enh | Enh | Enh |
| rs2594950 | Enh | Other | Enh |
| rs62172372 | Enh | Other | Other |
| rs2250390 | Enh | Other | Enh |
| rs736730 | Enh | Other | Other |
| rs7580831 | Enh | Other | Other |
| rs11886117 | Enh | Other | Other |
| rs568624328 | Enh | Other | Enh |
| rs62236869 | Enh | Enh | Other |
| rs16861312 | Enh | Enh | Enh |
| rs2284837 | Enh | Enh | Enh |
| rs116791817 | Enh | Enh | Other |
| rs145013093 | Enh | Enh | Other |
| rs114463740 | Enh | Enh | Other |
| rs35704688 | Enh | Other | Other |
| rs34915214 | Enh | Other | Other |
| rs189008 | Enh | Other | Other |
| rs992018 | Enh | Enh | Enh |
| rs443387 | Enh | Other | Enh |
| rs430130 | Enh | Other | Enh |
| rs2617407 | Enh | Other | Other |
| rs6862748 | Enh | Other | Other |
| rs12234019 | Enh | Enh | Enh |
| rs2546985 | Enh | Other | Other |
| rs7732646 | Enh | Other | Other |
| rs145214406 | Enh | Other | Enh |
| rs2517482 | Enh | Other | Other |
| rs2535316 | Enh | Enh | Other |
| rs111844274 | Enh | Other | Other |
| rs10948226 | Prom | Other | Enh/prom |
| rs56253836 | Enh | Enh | Enh |
| rs56178742 | Enh | Enh | Enh |
| rs7749409 | Enh | Enh | Other |
| rs113880644 | Enh | Other | Other |
| rs61543455 | Enh | Enh | Other |
| rs3777446 | Enh | Other | Other |
| rs113628826 | Enh/prom | Other | Other |
| rs34044181 | Enh/prom | Enh/prom | Enh/prom |
| rs9355156 | Enh | Other | Other |
| rs28585071 | Enh | Other | Enh |
| rs73288499 | Enh | Other | Other |
| rs6962943 | Enh | Enh | Other |
| rs2892794 | Enh | Enh | Enh |
| rs6461209 | Enh | Other | Other |
| rs2215169 | Enh/prom | Other | Other |
| rs144198315 | Enh | Enh | Enh |
| rs73402679 | Enh/prom | Enh | Enh |
| rs10281213 | Enh | Enh | Other |
| rs1547960 | Enh | Other | Other |
| rs1547959 | Enh | Other | Other |
| rs2618481 | Enh | Other | Other |
| rs3808422 | Enh | Other | Enh |
| rs12544624 | Enh | Enh | Enh |
| rs6991351 | Enh | Other | Enh |
| rs6991756 | Enh | Other | Enh |
| rs16934621 | Enh | Other | Other |
| rs34946727 | Enh | Other | Other |
| rs10868771 | Enh | Other | Other |
| rs4330775 | Enh | Other | Enh |
| rs10868772 | Enh | Other | Other |
| rs73520181 | Enh | Other | Other |
| rs17662822 | Enh | Other | Enh |
| rs1159798 | Enh | Other | Enh |
| rs1896246 | Enh/prom | Enh/prom | Other |
| rs1896244 | Enh/prom | Enh/prom | Other |
| rs11376072 | Enh | Enh/prom | Other |
| rs1649043 | Enh | Other | Enh |
| rs1649038 | Enh | Enh | Other |
| rs7921371 | Enh | Enh | Other |
| rs7090615 | Enh | Enh | Enh |
| rs35202269 | Enh | Other | Other |
| rs10767630 | Enh | Other | Other |
| rs10835154 | Enh | Other | Other |
| rs7483162 | Enh | Other | Enh |
| rs79241864 | Enh | Enh | Other |
| rs618767 | Enh | Enh | Other |
| rs567064085 | Prom | Other | Other |
| rs616781 | Enh | Enh | Enh |
| rs549809 | Enh | Other | Enh |
| rs549941 | Enh | Other | Enh |
| rs550621 | Enh | Other | Enh |
| rs12303073 | Enh | Other | Enh |
| rs55643039 | Enh | Enh/prom | Enh |
| rs863750 | Enh | Enh | Enh |
| rs825452 | Enh | Enh | Enh |
| rs7296124 | Enh | Other | Other |
| rs1753633 | Enh/prom | Other | Enh/prom |
| rs770396 | Enh | Other | Other |
| rs9557081 | Enh | Enh | Other |
| rs6573316 | Enh | Enh | Enh |
| rs10136948 | Enh | Other | Other |
| rs17733282 | Enh | Enh | Enh |
| rs56259139 | Enh | Enh | Enh/prom |
| rs113099844 | Enh | Other | Other |
| rs112375829 | Enh | Enh | Other |
| rs78181938 | Enh | Enh | Other |
| rs10518707 | Enh | Other | Other |
| rs746978 | Enh | Other | Other |
| rs2541583 | Enh/prom | Enh | Other |
| rs350215 | Enh | Enh | Other |
| rs3841243 | Prom | Prom | Enh/prom |
| rs3169494 | Prom | Prom | Enh/prom |
| rs3812997 | Prom | Enh/prom | Prom |
| rs9940646 | Enh | Other | Other |
| rs79434985 | Enh | Enh | Enh |
| rs8066982 | Enh | Enh | Enh |
| rs2107565 | Enh | Other | Other |
| rs11653144 | Enh/prom | Prom | Other |
| rs6503475 | Enh | Other | Other |
| rs11279297 | Enh | Other | Other |
| rs535521250 | Enh | Enh | Enh |
| rs575478168 | Enh | Enh | Enh |
| rs7207503 | Enh | Enh | Other |
| rs1476917 | Enh | Enh | Other |
| rs9897571 | Enh | Enh | Enh |
| rs179903 | Prom | Other | Other |
| rs2292852 | Enh | Enh | Other |
| rs7222241 | Enh | Other | Other |
| rs7223613 | Enh | Other | Other |
| rs578203928 | Enh | Other | Other |
| rs6086318 | Enh | Enh | Other |
| rs6077302 | Enh | Enh | Other |
| rs6077991 | Enh | Enh | Enh |
| rs6088703 | Enh | Other | Enh/prom |
| rs6066132 | Enh | Other | Other |
| rs5764824 | Enh | Other | Other |
| rs66798233 | Enh | Other | Enh |

[a]Out of a total 57,235 BMD-associated index/proxy SNPs, this set of 154 unique EnhPro SNPs was determined. EnhPro SNPs were defined as BMD GWAS SNPs that preferentially overlapped strong enhancer (enh) or promoter (prom) chromatin and a narrow peak of histone H3 lysine K27 acetylation (H3K27ac) in osteoblasts (ostb) but in no more than three of the following 12 types of cell cultures according to the Roadmap database (Roadmap Hub, http://genome.ucsc.edu): skin fibroblasts, lung fibroblasts, fetal lung fibroblasts (IMR90), foreskin or dermal keratinocytes, melanocytes, astrocytes, umbilical cord endothelial cells, myoblasts, mammary epithelial cells, embryonic stem cells and a lymphoblastoid cell line, excluding mesenchymal stem cells (MSC) and chondrocytes (chond). In addition, EnhPro SNPs had to overlap a narrow-peak DNaseI hypersensitivity site (DHS) in ostb. NA, not applicable

[b]Tier-1 SNPs were EnhPro SNPs that are predicted to overlap allele-specific transcription factor binding sites related to bone biology. The genes associated with Tier-1 SNPs had to be preferentially expressed in ostb compared with 11 other cell types (Supplemental Table S7) or significantly associated with a skeletal system Gene Ontology term (Supplemental Table S5). In addition to the 13 index/proxy Tier-1 SNPs (shown in blue font), we identified three more Tier-1 SNPs (rs11006188 and rs1982173 associated with *BICC1* and rs80019710 associated with *HMGA2* by examining imputed eBMD and TB-BMD SNPs with $p < 6.6 \times 10^{-9}$ and $p < 5 \times 10^{-8}$, respectively (Morris JA et al. and Medina-Gomez C et al., Supplemental Table S1)

[c]For the three indicated SNPs (rs2504105, rs2504104 and rs2504103) associated with *DAAM2* , an exception was made to include them as Tier-1 SNPs even though four of the 12 heterologous cell types displayed a narrow peak of H3K27ac overlapping the SNP instead of no more than three, as was otherwise required

**Supplemental Table S4. Functional clustering analysis (DAVID[a]) of the 2,157 reference genes associated with BMD GWAS**

| Gene ontology (GO) terms significantly enriched among all GWAS SNP-associated genes[b] | Cluster enrichment score[c] | Reference gene count | Fold enrichment | *p* -value[d] |
|---|---|---|---|---|
| **Cluster 1** | 8.7 | | | |
| GO:0001501~skeletal system development | | 65 | 3.2 | 1.06E-16 |
| GO:0060348~bone development | | 28 | 3.5 | 1.30E-08 |
| GO:0001503~ossification | | 25 | 3.4 | 2.27E-07 |
| GO:0001649~osteoblast differentiation | | 12 | 4.4 | 4.89E-05 |
| **Cluster 2** | 8.5 | | | |
| GO:0048598~embryonic morphogenesis | | 53 | 2.7 | 9.41E-11 |
| GO:0060173~limb development | | 27 | 4.1 | 1.01E-09 |
| GO:0048736~appendage development | | 27 | 4.1 | 1.01E-09 |
| GO:0035107~appendage morphogenesis | | 26 | 4.1 | 2.11E-09 |
| GO:0035108~limb morphogenesis | | 26 | 4.1 | 2.11E-09 |
| GO:0035113~embryonic appendage morphogenesis | | 22 | 3.9 | 9.49E-08 |
| GO:0030326~embryonic limb morphogenesis | | 22 | 3.9 | 9.49E-08 |
| **Cluster 3** | 6.7 | | | |
| GO:0003700~transcription factor activity | | 114 | 1.8 | 4.83E-10 |
| GO:0030528~transcription regulator activity | | 157 | 1.6 | 1.07E-09 |
| GO:0051252~regulation of RNA metabolic process | | 174 | 1.5 | 2.17E-08 |
| GO:0006355~regulation of transcription, DNA-dependent | | 169 | 1.5 | 6.17E-08 |
| GO:0045449~regulation of transcription | | 230 | 1.4 | 6.94E-08 |
| dna-binding | | 167 | 1.5 | 1.71E-07 |
| transcription regulation | | 177 | 1.4 | 3.22E-07 |
| Transcription | | 179 | 1.4 | 5.43E-07 |
| GO:0003677~DNA binding | | 201 | 1.3 | 7.05E-06 |
| nucleus | | 315 | 1.2 | 4.96E-05 |
| GO:0006350~transcription | | 176 | 1.3 | 1.05E-04 |
| **Cluster 4** | 6.5 | | | |
| GO:0007389~pattern specification process | | 45 | 2.6 | 6.41E-09 |
| GO:0003002~regionalization | | 36 | 2.8 | 3.33E-08 |
| GO:0009953~dorsal/ventral pattern formation | | 17 | 4.4 | 7.53E-07 |
| GO:0009952~anterior/posterior pattern formation | | 23 | 2.6 | 8.23E-05 |
| **Cluster 5** | 6.1 | | | |
| wnt signaling pathway | | 26 | 3.5 | 6.03E-08 |
| GO:0016055~Wnt receptor signaling pathway | | 27 | 3.2 | 2.82E-07 |
| hsa05217:Basal cell carcinoma | | 17 | 4.5 | 4.17E-07 |
| hsa04310:Wnt signaling pathway | | 25 | 2.4 | 7.44E-05 |

[a]Database for Annotation, Visualization and Integrated Discovery (DAVID, v6.7) was used to look for significant enrichment among the genes in Supplemental Table S2: https://david-d.ncifcrf.gov/

[b]Only the top 5 clusters are presented here

[c]The overall enrichment score for the cluster based on the modified one-tail Fisher Exact p-value (EASE score) of each term members

[d]EASE score of each GO term

**Supplemental Table S5. Analysis of 31 EnhPro SNPs associated with skeleton-related genes by GREAT analysis of 154 EnhPro SNPs[a]**

| | Expression ratio of ostb, MSC or chond to the median RPKM of 11 other heterologous cell cultures[c] | | | Average RPKM of technical duplicates from ENCODE | | | | | |
|---|---|---|---|---|---|---|---|---|---|
| Gene[b] | Ostb/non-ostb (excl. MSC & chond)[d] | MSC/non-MSC (excl. ostb & chond) | Chond/non-chond (excl. ostb & MSC) | Ostb | MSC | Chond | EnhPro SNP (distance to gene's TSS) | Overlapping allele-specific TFBS[d] | Comments |
| HMGA2 | 0.30 | 2.43 | 0.25 | 0.53 | 4.20 | 0.44 | rs17101510 (-230716) | Yes | One of the 16 Tier 1 SNPs |
| " | " | " | " | " | " | " | rs12296417 (-230185) | Yes | " |
| NPR3 | 14.61 | 7.21 | 22.31 | 3.17 | 1.57 | 4.85 | rs1173771 (+103488) | Yes | One of the 16 Tier 1 SNPs chosen partly for its gene's ostb-preferential expression |
| " | " | " | " | " | " | " | rs7733331 (+117306) | Yes | " |
| TBX15 | 296.80 | 410.35 | 479.88 | 2.07 | 2.86 | 3.34 | rs6680737 (+10548) | Yes | Previously examined by the current authors (Zhang, X et al., 2020, PMID: 31975641 ) |
| BMP5 | 35.08 | 267.87 | 4.32 | 0.02 | 0.19 | 0.00 | rs7749409 (-165644) | No | No allelic TFBS |
| DLX6 | #DIV/0! | #DIV/0! | #DIV/0! | 0.07 | 0.10 | 0.07 | rs73402679 (-42887) | No | No allelic TFBS |
| SIX1 | 23.58 | 30.43 | 76.56 | 0.98 | 1.26 | 3.17 | rs6573316 (+58151) | No | No allelic TFBS |
| SMAD3 | 1.43 | 1.96 | 3.30 | 2.02 | 2.76 | 4.64 | rs113099844 (-233197) | No | No allelic TFBS |
| " | " | " | " | " | " | " | rs112375829 (-233110) | No | No allelic TFBS |
| " | " | " | " | " | " | " | rs78181938 (-233070) | No | No allelic TFBS |
| " | " | " | " | " | " | " | rs10518707 (+7439) | No | No allelic TFBS |
| " | " | " | " | " | " | " | rs746978 (+129881) | No | No allelic TFBS |
| TRPS1 | 1.64 | 3.45 | 3.92 | 4.20 | 8.84 | 10.04 | rs3808422 (+164375) | No | No allelic TFBS |
| WNT7B | 6.83 | 74.75 | 2.41 | 0.01 | 0.12 | 0.00 | rs5764824 (+293794) | No | No allelic TFBS |
| RUNX2 | 6.07 | 4.14 | 0.21 | 1.15 | 0.78 | 0.04 | rs111844274 (-589566) | No | No allelic TFBS |
| " | " | " | " | " | " | " | rs10948226 (+3068) | No | No allelic TFBS |
| PDGFC | 2.15 | 2.36 | 1.26 | 5.42 | 5.94 | 3.17 | rs34915214 (+376271) | No | No allelic TFBS |
| " | " | " | " | " | " | " | rs35704688 (+376374) | No | No allelic TFBS |
| NOV | 1.22 | 3.53 | 4.16 | 0.40 | 1.15 | 1.36 | rs12544624 (+98583) | NE | Documentation of its gene's relationship to bone biology in the literature but only very low-to-low expression in ostb, chond, or MSC cell cultures |
| " | " | " | " | " | " | " | rs6991351 (+128420) | NE | " |
| " | " | " | " | " | " | " | rs6991756 (+128421) | NE | " |
| FLI1 | 1.07 | 0.49 | 1.07 | 0.58 | 0.27 | 0.58 | rs616781 (+121450) | NE | Documentation of its gene's relationship to bone biology in the literature but only very low expression in ostb, chond, or MSC cell cultures |
| " | " | " | " | " | " | " | rs549809 (+121652) | NE | " |
| " | " | " | " | " | " | " | rs549941 (+121689) | NE | " |
| " | " | " | " | " | " | " | rs550621 (+121697) | NE | " |
| TRIP11 | 1.06 | 1.19 | 1.24 | 2.80 | 3.14 | 3.28 | rs17733282 (+59840) | No | Documentation of its gene's relationship to bone biology in the literature but only very low expression in ostb, chond, or MSC cell cultures |
| NOG | 0.24 | 0.42 | 0.04 | 0.02 | 0.03 | 0.00 | rs7207503 (-452410) | NE | Documentation of its gene's relationship to bone biology in the literature but negligible expression in ostb, MSC, & chond cell cultues |
| " | " | " | " | " | " | " | rs1476917 (-451382) | NE | " |
| " | " | " | " | " | " | " | rs9897571 (-422311) | NE | " |
| C5orf20 | #DIV/0! | #DIV/0! | #DIV/0! | 0.00 | 0.00 | 0.00 | rs6862748 (+8854) | NE | Negligible expression in ostb, MSC, & chond. |

[a]154 EnhPro SNPs were analyzed for significant enrichment skeleton-related terms using the GREAT v4.0.4 (Genomic Regions Enrichment of Annotations Tool; http://great.stanford.edu/) with default association rule settings

[b]16 genes associated with skeletal system developmental (GO: 0001501; binomial FDR Q-value = $8 \times 10^{-6}$) were obtained from GREAT analysis; the eight genes most strongly associated with ostb, chond, or MSC are shown in blue font; the third *HMGA2* Tier-1 SNP rs80019710 is not listed because it was obtained as an imputed SNPs (Table 1)

[c]Expression ratios were calculated as described in Supplemental Table S2; ostb, osteoblasts; MSC, bone marrow-derived mesenchymal stem cells; chond, chondrocytes; fib, fibroblasts; excl, excluding [d]Manual curation of predicted TFBS from TRANSFAC v2019.3 (see Supplemental Materials and Methods); NE, not examined for predicted allele-specific TFBS

**Supplemental Table S6. The transcription factors predicted to bind with allele specificity to the 16 Tier-1 SNP-containing sequences are known to be associated with the skeletal system[a]**

| Gene | Tier-1 SNP[b] | Allele with predicted allele-specific TF binding | Name of TF or TF family | PWM Id[c] | 21-base DNA sequence centered on the SNP (underlined) | Brief characterizations of the transcription factors (TFs) for the allelic transcription factor binding sites (TFBS) at Tier-1 SNPs: Representative references (first author, year, PMID) | |
|---|---|---|---|---|---|---|---|
| | | | | | | Repressor (Rep) or activator (Act) activity of the TF | Associations of the TFs with the skeletal system |
| BICC1 | rs112597538 | Ref | SREBP1 | V$SREBF1_07 | gggtggcggGTGGCGTGgGcg | Rep or Act: GeneCards[d]; Ide T, 2004, PMID: 15048126 | SREBP1, important for the mineralization of osteoblast (ostb) cultures in vitro (Gorski JP, 2011, PMID: 21075843; Medina-Gomez C, 2017, PMID: 28743860); SREBP-1c upregulates MAFB, which is required for proper osteoclast proliferation, differentiation and activity in osteoclast progenitors of mice (Menendez-Gutierrez MP, 2015, PMID: 25574839) |
| " | " | Ref | SREBP2 | V$SREBP2_Q6 | gggtgGCggGTGGCgtgggcg | Rep or Act: GeneCards; Ide T, 2004, PMID: 15048126 | In the absence of MSX1, SREBP2 can suppress ostb differentiation in human dental pulp stem cells (Goto N, 2016, PMID: 27648077); SREBP2 is required for osteoclastogenesis (Zheng ZG, 2020,PMID: 31907393; Inoue K, 2015, PMID: 26319416) |
| " | rs1896245 | Ref | SATB1 | V$SATB1_Q5_01 | tattagtagATAAAttcttct | Rep or Act: GeneCards; Mir R, 2012, PMID: 22998183 | SATB1 promotes osteoclastogeneisis (Li J, 2017, PMID: 29442043); homozygous knockout resulted in a decrease in body size and in postnatal growth retardation (http://www.informatics.jax.org/allele/MGI:2386676; http://www.informatics.jax.org/allele/MGI:4436287); heterozygotes have increased bone mineral content in both female and male mice (http://www.informatics.jax.org/allele/MGI:5548358). SATB1 is closely related to SATB2, which regulates the proliferation of pre-ostb, may interact with SATB1, and, like SATB2, regulates higher-order chromatin organization (Dowry T, 2019, PMID: 31325654; Zhou LQ, 2012, PMID: 22825848; Naik R, 2019, PMID: 30413763) |
| " | rs1896243 | Alt | TCF3 | V$TCF3_Q6 | attactaCTTTGAaacaacca | Rep or Act: GeneCards | TCF3, a key transcription factor of canonical Wnt pathway, promotes osteogenic differentiation both *in vitro* and *in vivo* (Liu W, 2013, PMID: 23492770) |
| " | rs1658429 | Alt | AP-1 | V$AP1_C | cataatatatCTGAGTCAgca | Act: GeneCards; Bejjani F, 2019, PMID: 31034924 | AP-1 may affect enhancer selection (Madrigal P, 2018, PMID: 29778529); however, the effects of AP-1 on osteogenesis depend on the exact JUN:FOS heterodimers/homodimers (McCabe LR, 1996, PMID: 8828501; Eferl R, 2004, PMID: 15229648; Xia B, 2017, PMID: 29203636) |
| " | rs11006188 | Ref | SMAD | V$SMAD_Q4 | ggtgcctgCTGTCTGtcctta | Rep or Act: GeneCards; Uchiyama Y, 2008, PMID: 18466073 | SMAD family proteins play important roles in osteogenesis and chondrogenesis (Song B, 2009, PMID: 19926329); SMAD3, an intracellular signal transducer and TF, regulates osteogenesis and chondrogenesis and inhibits early healing of bone fractures and is activated by TGFB1, a critical mediator of bone marrow MSC recruitment to form ostb in bone repair (GeneCards); SMAD3 inhibits bone resorption (via the RANKL) when it is not monoubiquitinated (Xu Z, 2017, PMID: 28216630; Lin HT, 2018, PMID: 30483761; Liu W, 2018, PMID: 30482881; Xie Y, 2018, PMID: 30024590; Nguyen J, 2013, PMID: 23308287) |
| " | " | Ref | GLIS3 | V$ZNF515_Q6 | ggtgCCTGCTGTctgtcctta | Rep or act: GeneCards | Glis3 is highly expressed in human ostb and acts synergistically with BMP2 and Shh in enhancing ostb differentiation in multipotent C3H10T1/2 cells (Beak JY, 2007, PMID: 17488195) |
| " | " | Alt | NR2F6 | V$EAR2_Q2 | ggtgcctGctCTcTGtCCTTa | Rep: GeneCards | Homozygous knockout of NR2F6 in both male and female mice exhibited abnormal bone structure and decreased bone mineral content (http://www.informatics.jax.org/allele/MGI:6153804) |
| " | rs1982173 | Ref | RBPJ | V$RBPJK_02 | tcttctGTGTGAGAAtcagat | Rep or act: GeneCards | Deletion of RBPJ in murine bone marrow MSC caused a dramatic high-bone-mass phenotype, diminished bone marrow MSC pool, and caused a rapid age-dependent bone loss (Tu XL, 2012, PMID: 22457635); Rbpj is a potent positive regulator of ostb differentiation and maturation in pluripotent mesenchymal Kusa-A1 cells (Wang SC, 2010, PMID: 20691157); RBPJ act as negative regulator of osteoclastogenesis in murine myeloid osteoclast lineage (Zhao BH, 2012, PMID: 22249448); Deletion of RBPJ resulted decrease in trabecular bone mass and increase in osteoclasts in both murine bone marrow and macrophages (Ma J, 2013, PMID: 23224519) |
| " | rs4948523 | Ref | PRDX2 | V$PRX2_Q2 | tacgtaagTaAAATTAAatat | Unknown | Prx II KO mice had a higher bone mass than did WT mice (Kim KM, 2019, PMID: 31160554); Prx II KO mice with lipopolysaccharide-induced, through JNK and STAT3, showed increased osteoclastogenesis compared with WT mice (ParkH, 2015, PMID: 25074339) |
| NPR3 | rs1173771 | Ref | SATB1 | V$SATB1_Q5_01 | ttgtgaATAAAttggtatgag | Rep or Act: GeneCards; Mir R, 2012, PMID: 22998183 | SATB1 promotes osteoclastogeneisis (Li J, 2017, PMID: 29442043); homozygous knockout had decreased body size and postnatal growth retardation (http://www.informatics.jax.org/allele/MGI:2386676; http://www.informatics.jax.org/allele/MGI:4436287); heterozygotes have increased in bone mineral content in both female and male mice (http://www.informatics.jax.org/allele/MGI:5548358). SATB1 is closely related to SATB2, which regulates the proliferation of pre-ostb, may interact with SATB1, and like SATB2 regulates higher-order chromatin organization (Dowry T, 2019, PMID: 31325654; Zhou LQ, 2012, PMID: 22825848; Naik R, 2019, PMID: 30413763) |
| " | rs7733331 | Ref | GTF2I | V$TFIII_Q6_01 | gtacAGGAtGTGaatatttgg | Rep and Act: GeneCards; Bu Y, 2011, PMID: 20568114 | Heterozygous deletion of the Gtf2i gene in mice gives craniofacial and general osteogenic defects (Enkhmandakh B, 2009, PMID: 19109438); GTF2I can regulate osteogenesis genes (Lazebnik MB, 2009, PMID: 19880526; Hakelien AM, 2014, PMID: 24898411) |
| LGR4 | rs10835153 | Alt | BPTF | V$FAC1_01 | gatctTGTTGTGtgaaaattc | Co-Act: Ma YQ, 2015, PMID: 26041917 | Abnormal phenotypes associated with BPTF in humans include short stature (HP: 0004322, https://hpo.jax.org/) and abnormalities in the skeletal system (HP:0000924, https://hpo.jax.org/) as is seen in mice models (http://www.informatics.jax.org/allele/MGI:6257533; http://www.informatics.jax.org/allele/MGI:3823037; http://www.informatics.jax.org/allele/MGI:3841298). |

| | | | | | | | |
|---|---|---|---|---|---|---|---|
| HMGA2 | rs80019710 | Alt | HMGA1 | V$HMGIY_Q3 | tcttccTTcATTTTtTTtttg | Rep or Act: GeneCards; Kim J, 1995, PMID: 7705411 | HMGA1 can up-regulate Wnt signaling and accelarates fracture healing (Xian L, 2017, PMID: 28452345; Zhang W, 2020, PMID: 31782345) |
| " | rs17101510 | Alt | YY1 | V$YY1_Q6_02 | tTaGCCATtTTgtacacatta | Rep or act: GeneCards | YY1 is implicated in osteoclast differenitiation and bone remodeling (Teitelbaum SL, 2000, PMID: 10968780; Shi ZQ, 2004, PMID: 15563837); YY1-cofactor complexes may act as either negative or positive regulators of ostb differentiation (Jeong HM, 2014, PMID: 24325869; Chen YH, 2018, PMID: 29637005) |
| " | rs12296417 | Alt | SATB1 | V$SATB1_Q5_01 | tatctgtATAAAtgtataatg | Rep or Act: GeneCards; Mir R, 2012, PMID: 22998183 | See the description for SATB1 for rs1896245 above |
| " | " | Alt | ZBTB20[e] | V$ZBTB20_03 | tatctgtataAATGTATAatg | Rep or Act: Nagao M, 2016, PMID: 27000654; Zhang H, 2018, PMID: 29700307 | Knockout of *Zbtb20* in chond results in delayed endochondral ossification and in postnatal growth retardation (Zhou GD, 2015, PMID: 25564625) |
| DAAM2 | rs2504105 | Ref | SMAD[e] | V$SMAD2_Q6, V$SMAD3_02, V$SMAD4_05 | tctggtcTCTAGACAgaacag | Rep or Act: GeneCards; Uchiyama Y, 2008, PMID: 18466073 | See the description for SMAD family proteins for rs11006188 above |
| " | rs2504104 | Alt | RUNX1 | V$AML1_01 | tgttctcTGTGGTtgcctggt | Rep or Act: Behrens K, 2016, PMID: 27076172 | RUNX1 promotes osteogenesis in bone marrow-derived mesenchymal stem cells (Luo Y, 2019, PMID: 30391794); *Runx1* expression negatively regulates osteoclastogenesis and osteoclast activity (Soung DY, 2014, PMID: 24606124; Paglia DN, 2019, PMID: 31769548) |
| " | " | Alt | RUNX2 | V$RUNX2_06 | tgttctCTGTGGTTgcctggt | Rep or Act: Jensen ED, 2007, PMID: 17725488 | RUNX2 is essential for ostb differentiation and chondrocyte maturation (Allas L, 2019, PMID: 30296494; Qin X, 2015, PMID: 25262822; Kundu M, 2002, PMID: 12434156; Soung DY, 2014, PMID: 24606124; Luo Y, 2019, PMID: 30391794; Komori T, 2019, PMID: 30987410) |
| " | " | Alt | CBFB | V$PEBP2B_Q6 | tgttctcTGTGGTtgcctggt | Rep or Act: GeneCards | CBFB regulates bone developoment by stabilizing Runx family proteins (Qin X, 2015, PMID: 25262822; Kundu M, 2002, PMID: 12434156) |
| " | rs2504103 | Alt | NFIC | V$NF1C_Q6 | tctgtagTTGGCtggtatgaa | Rep or Act: GeneCards; Ouellet S, 2006, PMID: 17130157 | NFIC plays important role in tooth and bone development because Nfic-deficient mice show abnormal tooth and bone formation (Roh SY, 2017, PMID: 28462188) |

[a]TRANSFAC v2019.3 (http://genexplain.com/transfac/) transcription factor binding site (TFBS) database was used to find candidates for predicted allele-specific TFBS; then the results were hand-curated (see Materials and Methods and Supplemental Methods) using the corresponding position weight matrixes (PWMs); only the results that passed hand curation are shown

[b]16 best candidate SNPs for regulatory osteoporosis-risk SNPs (Tier-1 SNPs) were determined from BMD GWAS SNPs that met epigenetic criteria, expression criteria for their associated gene, as well as overlap of the SNP with predicted allele-specific TFBS in osteoblasts (ostb) as described in the main text

[c]PWM ID or example of group PWM IDs for TF family from the TRANSFAC database that this analysis predicts to be an allele-specific TFBS at the DNA sequences containing the Tier-1 SNP; capitalized bases, highly conserved positions for the TF; underlined base, SNP position [d]GeneCards: http://www.genecards.org/

[e]For the Alt allele at Tier-1 SNP rs12296417, the predicted TFBS of ZBTB20 was accepted with one mismatch because the mismatched base is in a less conserved part of ZBTB20 PWM and the other eight bases of the PWM perfectly matched; SMAD2 and SMAD4 were predicted to perfectly match the DNA sequence around the Ref allele of Tier-1 SNP rs2504105, whereas the ref allele has one miscmatch out of eight to the SMAD3 PWM

**Supplementary Table S7. Most genes in the 1-Mb neighborhood of the five Tier-1 SNP-associated genes do not show the preferential expression in osteoblasts, mesenchymal stem cells, or chondrocytes that was seen in the Tier-1 SNP-associated genes[a]**

| Gene associated with Tier-1 SNP (red) & its neighbors (blue) | Expression ratio of ostb, MSC or chond to the median RPKM of 11 other heterologous cell cultures[b] | | | Average RPKM of technical duplicates from ENCODE | | | | | | | | | | | | | |
|---|---|---|---|---|---|---|---|---|---|---|---|---|---|---|---|---|---|
| | Ostb/non-ostb (excl. MSC & chond) | MSC/non-MSC (excl. ostb & chond) | Chond/non-chond (excl. ostb & MSC) | Ostb | MSC | Chond | Aortic endothelial cells | Follicle dermal papilla cells | HMEC | Placental pericytes | Saphous vein endothelial cells | White preadipocytes | Fetal lung fib (IMR90) | Diploid skin fib | Epidermal melanocytes (foreskin) | Epidermal melanocytes (cheek/temple) | Myoblasts |
| BICC1 | 5.3 | 4.7 | 5.3 | 7.1 | 6.3 | 7.1 | 0.1 | 6.7 | 1.3 | 1.5 | 0.0 | 1.7 | 2.6 | 2.7 | 0.2 | 0.2 | 1.4 |
| TFAM | 0.6 | 0.7 | 0.9 | 0.7 | 0.8 | 1.0 | 2.8 | 0.5 | 1.0 | 0.8 | 1.6 | 1.2 | 1.0 | 0.6 | 1.7 | 0.8 | 1.9 |
| FAM133CP | 7.3 | 2.3 | 9.0 | 0.5 | 0.2 | 0.6 | 0.0 | 0.3 | 0.1 | 0.1 | 0.0 | 0.1 | 0.0 | 0.2 | 0.1 | 0.0 | 0.1 |
| PHYHIPL | #DIV/0! | #DIV/0! | #DIV/0! | 0.0 | 0.0 | 0.0 | 0.0 | 0.0 | 0.0 | 0.0 | 0.0 | 0.0 | 0.0 | 0.0 | 0.0 | 0.0 | 0.0 |
| FAM13C | 12.2 | 9.6 | 12.4 | 0.3 | 0.2 | 0.3 | 0.0 | 0.0 | 0.1 | 0.0 | 0.4 | 0.0 | 0.2 | 0.1 | 0.0 | 0.0 | 0.0 |
| LINC00844 | #DIV/0! | #DIV/0! | #DIV/0! | 0.0 | 0.0 | 0.0 | 0.0 | 0.0 | 0.0 | 0.0 | 0.0 | 0.0 | 0.0 | 0.0 | 0.0 | 0.0 | 0.0 |
| CCEPR | #VALUE! | #VALUE! | #VALUE! | NA[c] | NA | NA | NA | NA | NA | NA | NA | NA | NA | NA | NA | NA | NA |
| NPR3 | 14.6 | 7.2 | 22.3 | 3.2 | 1.6 | 4.8 | 0.0 | 0.9 | 0.2 | 0.1 | 0.1 | 0.7 | 4.6 | 1.0 | 0.0 | 0.0 | 2.6 |
| MTMR12 | 0.6 | 1.0 | 0.9 | 0.8 | 1.3 | 1.1 | 2.7 | 1.1 | 2.1 | 1.2 | 2.6 | 1.2 | 1.8 | 1.0 | 1.3 | 0.8 | 1.3 |
| ZFR | 0.8 | 1.0 | 0.8 | 6.8 | 8.8 | 6.7 | 7.5 | 11.0 | 8.6 | 9.4 | 10.4 | 7.5 | 17.3 | 7.2 | 9.0 | 7.8 | 6.4 |
| SUB1 | 0.8 | 0.9 | 0.9 | 4.5 | 5.3 | 5.4 | 6.3 | 5.7 | 4.2 | 5.5 | 4.8 | 8.8 | 5.1 | 6.5 | 6.0 | 3.5 | 10.9 |
| LINC02120 | #DIV/0! | #DIV/0! | #DIV/0! | 0.0 | 0.0 | 0.0 | 0.0 | 0.0 | 0.0 | 0.0 | 0.0 | 0.0 | 0.0 | 0.0 | 0.0 | 0.0 | 0.0 |
| MIR579 | #VALUE! | #VALUE! | #VALUE! | NA | NA | NA | NA | NA | NA | NA | NA | NA | NA | NA | NA | NA | NA |
| LGR4 | 5.1 | 3.4 | 2.2 | 3.0 | 2.0 | 1.3 | 0.5 | 2.2 | 2.3 | 0.6 | 0.2 | 0.2 | 0.6 | 1.4 | 0.9 | 0.8 | 0.3 |
| LGR4-AS1 | #DIV/0! | #DIV/0! | #DIV/0! | 0.0 | 0.0 | 0.0 | 0.0 | 0.0 | 0.0 | 0.0 | 0.0 | 0.0 | 0.0 | 0.0 | 0.0 | 0.0 | 0.0 |
| CCDC34 | 1.0 | 1.5 | 1.1 | 0.1 | 0.2 | 0.2 | 0.2 | 0.1 | 0.1 | 0.1 | 0.1 | 0.2 | 0.2 | 0.1 | 0.3 | 0.1 | 0.2 |
| FIBIN | 2.5 | 1.0 | 1.0 | 0.4 | 0.2 | 0.2 | 0.0 | 3.2 | 0.3 | 2.8 | 0.0 | 0.1 | 0.2 | 0.4 | 0.0 | 0.0 | 0.2 |
| BBOX1 | #DIV/0! | #DIV/0! | #DIV/0! | 0.0 | 0.0 | 0.0 | 0.0 | 0.0 | 0.1 | 0.0 | 0.0 | 0.0 | 0.0 | 0.0 | 0.0 | 0.0 | 0.0 |
| BBOX1-AS1 | #DIV/0! | #DIV/0! | #DIV/0! | 0.1 | 0.0 | 0.0 | 0.0 | 0.0 | 0.0 | 0.1 | 0.0 | 0.0 | 0.0 | 0.0 | 0.0 | 0.0 | 0.0 |
| LIN7C | 1.0 | 1.5 | 1.5 | 2.6 | 4.2 | 4.0 | 3.2 | 2.7 | 4.0 | 2.7 | 2.5 | 2.3 | 3.1 | 2.4 | 3.1 | 1.8 | 3.5 |
| BDNF-AS | 0.8 | 1.7 | 2.4 | 0.1 | 0.1 | 0.2 | 0.1 | 0.1 | 0.1 | 0.2 | 0.0 | 0.1 | 0.0 | 0.1 | 0.1 | 0.1 | 0.1 |
| BDNF | 1.9 | 4.0 | 1.6 | 0.4 | 0.9 | 0.4 | 0.0 | 1.5 | 0.2 | 3.2 | 0.1 | 0.0 | 18.5 | 0.3 | 0.2 | 0.3 | 0.1 |
| LINC00678 | #DIV/0! | #DIV/0! | #DIV/0! | 0.0 | 0.0 | 0.0 | 0.0 | 0.0 | 0.0 | 0.0 | 0.0 | 0.0 | 0.0 | 0.0 | 0.0 | 0.0 | 0.0 |
| MIR8087 | #VALUE! | #VALUE! | #VALUE! | NA | NA | NA | NA | NA | NA | NA | NA | NA | NA | NA | NA | NA | NA |
| HMGA2 | 0.3 | 2.4 | 0.3 | 0.5 | 4.2 | 0.4 | 2.7 | 0.5 | 0.9 | 13.2 | 1.7 | 2.9 | 7.8 | 0.6[d] | 0.0 | 0.0 | 3.7 |
| LINC02454 | 0.7 | 2.4 | 0.2 | 0.2 | 0.8 | 0.1 | 0.5 | 0.1 | 0.5 | 0.2 | 0.5 | 0.4 | 0.3 | 0.2[d] | 0.2 | 0.2 | 0.5 |
| RPSAP52 | 0.5 | 3.4 | 1.0 | 0.1 | 0.8 | 0.2 | 0.4 | 0.1 | 0.1 | 0.3 | 0.1 | 0.8 | 0.4 | 0.2 | 0.0 | 0.0 | 0.8 |
| HMGA2-AS1 | 27.0 | 12.7 | 22.3 | 0.4 | 0.2 | 0.3 | 0.0 | 0.1 | 0.0 | 0.0 | 0.0 | 0.2 | 0.0 | 0.8 | 0.0 | 0.0 | 0.1 |
| LEMD3 | 0.9 | 1.0 | 0.6 | 1.6 | 1.7 | 1.2 | 1.5 | 2.0 | 3.3 | 2.2 | 1.9 | 1.7 | 1.7 | 1.8 | 1.5 | 1.5 | 2.1 |
| MSRB3 | 1.6 | 0.7 | 1.3 | 3.2 | 1.3 | 2.6 | 2.0 | 3.1 | 3.5 | 2.7 | 2.4 | 0.8 | 2.3 | 1.9 | 0.3 | 0.6 | 1.1 |
| LOC100507065 | 0.1 | 9.6 | 0.1 | 0.0 | 0.1 | 0.0 | 0.0 | 0.0 | 0.0 | 0.0 | 0.0 | 3.0 | 0.0 | 0.0 | 0.0 | 0.0 | 1.4 |
| LLPH | 1.4 | 1.2 | 2.2 | 0.3 | 0.3 | 0.6 | 0.4 | 0.2 | 0.1 | 0.2 | 0.2 | 0.2 | 0.5 | 0.4 | 0.5 | 0.3 | 0.5 |
| TMBIM4 | 0.8 | 1.2 | 1.5 | 2.8 | 4.2 | 4.9 | 3.3 | 2.8 | 2.4 | 3.4 | 3.0 | 2.7 | 4.1 | 3.5 | 4.0 | 4.8 | 3.5 |
| LLPH-DT | 0.4 | 0.9 | 2.7 | 0.0 | 0.0 | 0.0 | 0.0 | 0.0 | 0.0 | 0.0 | 0.0 | 0.0 | 0.0 | 0.0 | 0.0 | 0.0 | 0.0 |
| LINC02425 | #DIV/0! | #DIV/0! | #DIV/0! | 0.0 | 0.0 | 0.0 | 0.0 | 0.0 | 0.0 | 0.0 | 0.0 | 0.0 | 0.0 | 0.0 | 0.0 | 0.0 | 0.0 |
| MIR6074 | #VALUE! | #VALUE! | #VALUE! | NA | NA | NA | NA | NA | NA | NA | NA | NA | NA | NA | NA | NA | NA |
| LOC105369187 | #VALUE! | #VALUE! | #VALUE! | NA | NA | NA | NA | NA | NA | NA | NA | NA | NA | NA | NA | NA | NA |
| DAAM2 | 20.6 | 6.8 | 34.6 | 1.2 | 0.4 | 2.1 | 0.0 | 3.6 | 0.0 | 0.0 | 0.0 | 0.0 | 0.8 | 2.3 | 3.6 | 2.7 | 0.1 |
| DAAM2-AS1 | #DIV/0! | #DIV/0! | #DIV/0! | 0.0 | 0.0 | 0.0 | 0.0 | 0.0 | 0.0 | 0.0 | 0.0 | 0.0 | 0.0 | 0.0 | 0.0 | 0.0 | 0.0 |
| KIF6 | 1.6 | 1.2 | 2.3 | 0.0 | 0.0 | 0.1 | 0.0 | 0.2 | 0.0 | 0.0 | 0.0 | 0.0 | 0.0 | 0.2 | 0.2 | 0.3 | 0.0 |
| MOCS1 | 1.4 | 2.2 | 1.3 | 1.2 | 1.8 | 1.1 | 0.8 | 0.8 | 0.3 | 1.2 | 0.7 | 0.9 | 0.9 | 0.8 | 0.8 | 1.0 | 0.8 |
| TDRG1 | #DIV/0! | #DIV/0! | #DIV/0! | 0.0 | 0.0 | 0.0 | 0.0 | 0.0 | 0.0 | 0.0 | 0.0 | 0.0 | 0.0 | 0.0 | 0.0 | 0.0 | 0.0 |
| LINC00951 | #DIV/0! | #DIV/0! | #DIV/0! | 0.0 | 0.0 | 0.0 | 0.0 | 0.0 | 0.0 | 0.0 | 0.0 | 0.0 | 0.0 | 0.0 | 0.0 | 0.0 | 0.0 |

[a]Expression levels, the average of duplicate RPKM (reads per kilobase million) determinations, except for human mammary epithlial cells (HMEC), for which only a single value was available

[b]Data from long RNA- seq (whole cell, total RNA > 0.2 kb, ENCODE/Cold Spring Harbor Lab; http://genome.ucsc.edu); expression ratios were calculated as described in Supplemental Table S2; ostb, osteoblasts; MSC, bone marrow-derived mesenchymal stem cells; chond, chondrocytes; HMEC, mammary epithelial cells; fib, fibroblasts; excl, excluding

[c]NA, not available

[d]These data are from a single primary skin fib cell culture; RNA-seq data from 504 independent primary skin fib cultures indicated specific transcription of *HMGA2* and *LINC02454* RNAs in skin fib vs. 50 human tissues and lymphoblastoid cell lines (Supplemental Fig. S8)

**Supplemental Table S8. Skeletal biology associations of nine genes associated with BMD GWAS-derived SNPs, a mouse limb phenotype and with negligible expression in a human osteoblast sample[a]**

| BMD GWAS assoc. reference genes[b] | Expression ratio of ostb, MSC or chond to the median RPKM of 11 other heterologous cell cultures[c] | | | Average RPKM of technical duplicates from ENCODE | | | Representative references for the gene's involvement in bone biology despite very low expression in the tested human osteoblast sample used by ENCODE[b] |
|---|---|---|---|---|---|---|---|
| | Ostb/non-ostb (excl. MSC & chond) | MSC/non-MSC (excl. ostb & chond) | Chond/non-chond (excl. ostb & MSC) | Ostb | MSC | Chond | |
| PKDCC | 0.18 | 0.09 | 1.10 | 0.09 | 0.04 | 0.56 | The Novel Protein Kinase Vlk Is Essential for Stromal Function of Mesenchymal Cells. Development. 2009;136(12):2069-2079, PMID: 19465597. Probst S, Zeller R, Zuniga A. The hedgehog target Vlk genetically interacts with Gli3 to regulate chondrocyte differentiation during mouse long bone development. Differentiation. 2013 Apr-Jun;85(4-5):121-30. PMID: 23792766. http://biogps.org/#goto=genereport&id=106522, appreciable expression of *Pkdcc* in mouse ostb. **Early chond or chond precursor cells might be the most important target of this gene.** |
| MEOX2 | 109.75 | 327.01 | 330.07 | 0.09 | 0.26 | 0.26 | Mankoo BS, Skuntz S, Harrigan I, et al. The concerted action of Meox homeobox genes is required upstream of genetic pathways essential for the formation, patterning and differentiation of somites. Development. 2003;130(19):4655-4664, PMID: 12925591 . Very low expression in human ostb but higher than non-bone related cell types and selectively expressed in mouse ostb vs. other cell types. http://biogps.org/#goto=genereport&id=17285, low expression in ostb. **Low expression in postnatal ostb, MSC, & chond may be due to this protein playing a key role mostly in somitogenesis and specifically sclerotome development.** |
| THRB | 0.27 | 1.99 | 0.94 | 0.04 | 0.31 | 0.15 | Bassett JH, Williams GR. Role of Thyroid Hormones in Skeletal Development and Bone Maintenance. Endocr Rev. 2016;37(2):135-187. PMID: 26862888. Cardoso LF, de Paula FJ, Maciel LM. Resistance to thyroid hormone due to mutations in the THRB gene impairs bone mass and affects calcium and phosphorus homeostasis. Bone. 2014;67:222-227, PMID: 25063548. http://biogps.org/#goto=genereport&id=21834, no signif expression detected in mouse ostb. **This protein affects bone through systemic regulation of calcium and phosphate metabolism.** |
| EN1 | 7.97 | 1.18 | 2.37 | 0.02 | 0.00 | 0.01 | Deckelbaum RA, Majithia A, Booker T, Henderson JE, Loomis CA. The homeoprotein engrailed 1 has pleiotropic functions in calvarial intramembranous bone formation and remodeling. Development. 2006 Jan;133(1):63-74, PMID: 16319118; http://biogps.org/#goto=genereport&id=13798, no detectable signal in mouse ostb or osteoclasts. **This homeobox protein may play its role mostly in pre-natal development & in osteoclasts** |
| CYP26B1 | 0.21 | 1.45 | 1.12 | 0.01 | 0.06 | 0.04 | Lind T, Sundqvist A, Hu L, Pejler G, Andersson G, Jacobson A, Melhus H. Vitamin a is a negative regulator of osteoblast mineralization. PLoS One. 2013 Dec 10;8(12):e82388, PMID: 24340023. http://biogps.org/#goto=genereport&id=232174, *Cyp26b1* expression in mouse ostb higher than in many other samples despite the negligible expression in the ENCODE ostb sample. **There might be technical reasons for these differences.** |
| GRIP1 | 0.55 | 1.41 | 1.44 | 0.01 | 0.02 | 0.02 | This match by GeneAnalytics of GRIP1 to mouse limb phenotype is not substantiated by any articles that we could find in PubMed for GRIP1, glutamate receptor interacting protein. However, there are such articles for NR3C1, glucocorticoid interacting proteinwhich was previously called GRIP1. http://biogps.org/#goto=genereport&id=74053, no significant expression of the glutamate receptor interacting protein-encoding gene in mouse ostb or osteoclasts but there is ostb-specific expression of *NR3C1* in human cells & selective expression in mice ostb & osteoclasts. **There may be some confusion in the Gene Ontology analysis due to the use of the same gene name for different genes.** |
| FGF18 | 3.29 | 14.89 | 32.97 | 0.00 | 0.02 | 0.04 | Nagayama T, Okuhara S, Ota MS, Tachikawa N, Kasugai S, Iseki S. FGF18 accelerates osteoblast differentiation by upregulating Bmp2 expression. Congenit Anom (Kyoto). 2013 Jun;53(2):83-8, PMID: 23751042. Liu Z, Lavine KJ, Hung IH, Ornitz DM. FGF18 is required for early chondrocyte proliferation, hypertrophy and vascular invasion of the growth plate. Dev Biol. 2007;302(1):80-91, PMID: 17014841. http://biogps.org/#goto=genereport&id=14172, there is specific expression of *FGF18* in mouse ostb. **FGF18 may act mostly at the early stages of chondrogenesis & endochondral ossification.** |
| RSPO2 | 0.32 | 84.81 | 163.33 | 0.00 | 0.78 | 1.51 | Zhu C, Zheng XF, Yang YH, Li B, Wang YR, Jiang SD, Jiang LS. LGR4 acts as a key receptor for R-spondin 2 to promote osteogenesis through Wnt signaling pathway. Cell Signal. 2016 Aug;28(8):989-1000. PMID: 27140682. http://biogps.org/#goto=genereport&id=239405, no significant expression of *Rspo2* in mouse ostb or osteoclasts. **RSPO2 is a secreted ligand that is preferentially expressed in chond and MSC and may exert its effects on bone largely by interacting with the LGR4 receptor on ostb; *LGR4* is one of the five Tier-1 SNP-associated genes (see main text).** |
| SP7 | #DIV/0! | #DIV/0! | #DIV/0! | 0.00 | 0.00 | 0.00 | Artigas N, Gámez B, Cubillos-Rojas M, Sánchez-de Diego C, Valer JA, Pons G, Rosa JL, Ventura F. p53 inhibits SP7/Osterix activity in the transcriptional program of osteoblast differentiation. Cell Death Differ. 2017 Dec;24(12):2022-2031, PMID: 28777372. Xing W, Godwin C, Pourteymoor S, Mohan S. Conditional disruption of the osterix gene in chondrocytes during early postnatal growth impairs secondary ossification in the mouse tibial epiphysis. Bone Res. 2019;7:24. Published 2019 Aug 5, PMID: 31646014. http://biogps.org/#goto=genereport&id=170574, mouse ostb, especially early in differentiation, specifically expressed SP7. **The gene encoding the widely studied protein osterix transcription factor is required for the conversion of pre-ostb to ostb and for transdifferentiation of chond to ostb during endochondral ossification. Importantly, there was negligible expression of *SP7* in human ostb or chond primary cells but chond derived by in vitro differentiation of MSC, displayed strong enhancer, promoter, and transcription-type chromatin not seen in other types of cell cultures (Supplemental Fig. S1).** |

[a]Out of 2,157 reference genes obtained from BMD GWAS (see Materials and Methods; Supplemental Table S2), 881 BMD GWAS-derived reference genes had negligible expression, namely, RPKM < 0.1 as the average of technical duplicates from a human osteoblast (ostb) sample that was part of the ENCODE project (http://www.genome.ucsc.edu/cgi-bin/hgTrackUi?hgsid=841267789_qpbJIFfHJRztPgHjcZVJek55ZfI9&c=chr8&g=wgEncodeCshlLongRnaSeq)

[b]Nine of the 881 genes with negligible expression in ostb were found to be associated with mouse limb phenotype (MP:0002109) by GeneAnalytics (http://geneanalytics.genecards.org/, score = 15.85); representative references for these genes being associated with ostb, chondrocyte (chond), or bone marrow-derived mesenchymal stem cells (MSC) are given as well as a link to the mouse microarray-based transcriptomic data summarizing expression microarray results for ostb, osteoclasts, and heterologous cell cultures and tissue from mice and an evaluation of the relevance of the gene to skeletal biology

[c]Expression ratios were determined as described for Supplemental Table S2
