## Supplemental Methods and Figs S1-S10 for "Prioritization of osteoporosis-associated GWAS SNPs using epigenomics and transcriptomics"

### Table of Contents

1. Supplementary Methods ........................................................................................2-3
2. List of Supplemental Tables S1-S8 ........................................................................4
3. Supplemental Figures S1- S10 ..............................................................................5-14
4. References .............................................................................................................15

### 1. Supplementary Methods

Total RNA-seq data (strand-specific; ENCODE/Cold Spring Harbor Lab) for 12 cell cultures types were examined rather than poly(A)$^+$ RNA-seq data (whole cell; ENCODE/Cold Spring Harbor lab; http://genome.ucsc.edu)[1, 2] because only the former ENCODE dataset includes osteoblasts (ostb), bone marrow-derived mesenchymal stem cells (MSC) and chondrocytes (chond). The 11 cell cultures that were compared to ostb for determining preferential transcription in ostb (RPKM ostb/median RPKM of non-ostb > 5) were as follows: dermal fibroblasts, fetal lung fibroblasts (IMR-90), myoblasts, human mammary epithelial cells (HMEC), foreskin or dermal melanocytes, aortic endothelial cells, follicle dermal papilla cells, pericytes, saphenous vein endothelial cells, and preadipocytes (https://genome.ucsc.edu/ENCODE/cellTypes.html); all but IMR-90 were primary cell cultures. The ostb had been obtained from the femoral bone of two individuals (62 yr male and 56 yr female); MSC from femoral bone marrow of two individuals (57 yr male and 60 yr female); and chond from knee or femoral cartilage of two individuals (64 yr male and 56 yr female) and supplied commercially (Promocell, each ostb, MSC, and chond lot had been tested for alkaline phosphatase/bone mineralization, CD44/CD105 CD31/CD45 plus ability to differentiate into ostb, chond, and adipocytes, or COL2A1 by immunohistochemistry, respectively, by the company). The same samples were used for CAGE analysis of 5' ends of RNA (cap analysis of gene expression, http://www.genome.ucsc.edu/cgi-bin/hgTrackUi?hgsid=841402551_77CNRn6hnLfRDA53aA0VH6iPxZhj&c=chr5&g=wgEncodeRikenCage).

For analysis of Roadmap chromatin segmentation data,[3] the following cell types were compared to ostb to determine SNP overlap of enhancer or promoter chromatin preferentially in ostb: dermal fibroblasts, fetal lung fibroblasts (IMR-90), myoblasts, human mammary epithelial cells (HMEC), foreskin melanocytes, skin epidermal or foreskin keratinocytes, astrocytes, umbilical cord endothelial cells, embryonic stem cells, post-natal lung fibroblasts, and a lymphoblastoid cell line (GM12878). All but IMR-90 and GM12878 are primary cells. The preferential enhancer or promoter chromatin designation required that ostb, but no more than three of these non-ostb cell types, could overlap strong enhancer or promoter chromatin, namely, states 1, 3, 8, or 9 from the Auxiliary 18-state Roadmap model (http://www.genome.ucsc.edu/cgi-bin/hgTrackUi?hgsid=841402551_77CNRn6hnLfRDA53aA0VH6iPxZhj&c=chr5&g=hub_24125_RoadmapConsolidatedAssaya27004). We used Roadmap DHS profiling (http://www.genome.ucsc.edu/cgi-bin/hgTrackUi?hgsid=841402551_77CNRn6hnLfRDA53aA0VH6iPxZhj&c=chr5&g=hub_24125_RoadmapConsolidatedAssaya23001) for many of the same samples. The ostb for both chromatin state segmentation and DHS were primary cells from a single donor of unknown gender or age (Roadmap code name E129)[3] with no further description. We found that the transcriptomic and chromatin state segmentation profiles for ostb markers indicated that the primary ostb cultures for both epigenomic and transcriptomic analyses were not pre-ostb but rather ostb that were transcribing genes associated with early, middle, or late stages of osteoblastogenesis (Supplemental Fig. S1). The MSC used for chromatin segmentation and DHS profiling[(3)] were from four donor bone marrow samples (Rikshopitalet University Hospital and supplied to the Broad Institute (https://www.ebi.ac.uk/vg/epirr/view/IHECRE00001042.1) although further characterization of these cells (Roadmap codename E026)[(3)] is not publicly available. Epigenetic profiles of chond (Roadmap codename E049)[(3)] used cells obtained by *in vitro* differentiation of the same MSC to chond. Chond samples used for epigenomics and transcriptomics expressed many of the ostb markers, as expected, but also expressed very much higher levels of *SOX5*, a marker of both early and hypertrophic chond, than the other cell types (Supplemental Figs. S1 and S2). There were some differences in deduced marker expression in the chond used for epigenomics and those for transcriptomics, which may be due to *in vitro* differentiated cells being studied for their epigenomics and primary cells for their transcriptomics. The epigenetic data for monocytes comes from $CD14^+$ blood cells (E121) in the Roadmap database.[(3)] Mammalian conserved sequences in Fig. 2 are from the UCSC Genome Browser's Element Conservation track using the placental mammal database.

For prediction of transcription factor binding sites (TFBS) using TRANSFAC (TRANSFAC Professional http://gene-regulation.com/, 2019.3/Vertebrate TFBS; matrix scores > 0.8, core scores > 0.9, Only High Quality Matrices, and Minimized False Positives ),[(4)] the input DNA was a 21-base sequence centered around the SNP allele. Manual curation of the matches obtained from TRANSFAC is as described in the main text. We also looked for evidence of *in vivo* binding to the DNA sequences containing the 16 Tier-1 SNPs (ChiP-seq; Factorbook 3; https://genome.ucsc.edu/cgi-bin/hgTrackUi?hgsid=841425465_BmeuanOYDg94aHdqmKmcRorxWIAL&c=chr1&g=encRegTfbsClustered) and UniBind; https://genome.ucsc.edu/cgi-bin/hgTrackUi?hgsid=841425869_cLZwutcxD1wbg8Y2dMlFFpZFz0kA&c=chr1&g=hub_1098543_UniBind) but identified none probably due to the very small number of transcription factor (TF) ChIP-seq profiles available for ostb.

### 2. List of Supplemental Tables (see Excel file)

**Supplemental Table S1.** The 38 BMD GWA studies used for this analysis

**Supplemental Table S2.** Expression ratios for some reference genes associated with BMD GWAS SNPs from the studies in Table S1: RNA-seq data for expression in osteoblasts, mesenchymal stem cells, or chondrocytes vs. in heterologous cell cultures

**Supplemental Table S3.** The 154 BMD GWAS SNPs that fit the epigenetic criteria to be EnhPro SNPs

**Supplemental Table S4.** Functional clustering analysis (DAVID) of the 2,157 reference genes associated with BMD GWAS

**Supplemental Table S5.** Analysis of 31 EnhPro SNPs associated with skeleton-related genes by GREAT analysis of 154 EnhPro SNPs

**Supplemental Table S6.** The transcription factors predicted to bind with allele specificity to the 16 Tier-1 SNP-containing sequences are known to be associated with the skeletal system

**Supplementary Table S7.** Most genes in the 1-Mb neighborhood of the five Tier-1 SNP-associated genes do not show the preferential expression in osteoblasts, mesenchymal stem cells, or chondrocytes that was seen in the Tier-1 SNP-associated genes

**Supplemental Table S8.** Skeletal biology associations of nine genes associated with BMD GWAS-derived SNPs, a mouse limb phenotype, and with negligible expression in a human osteoblast sample

### 3. Supplemental Figures

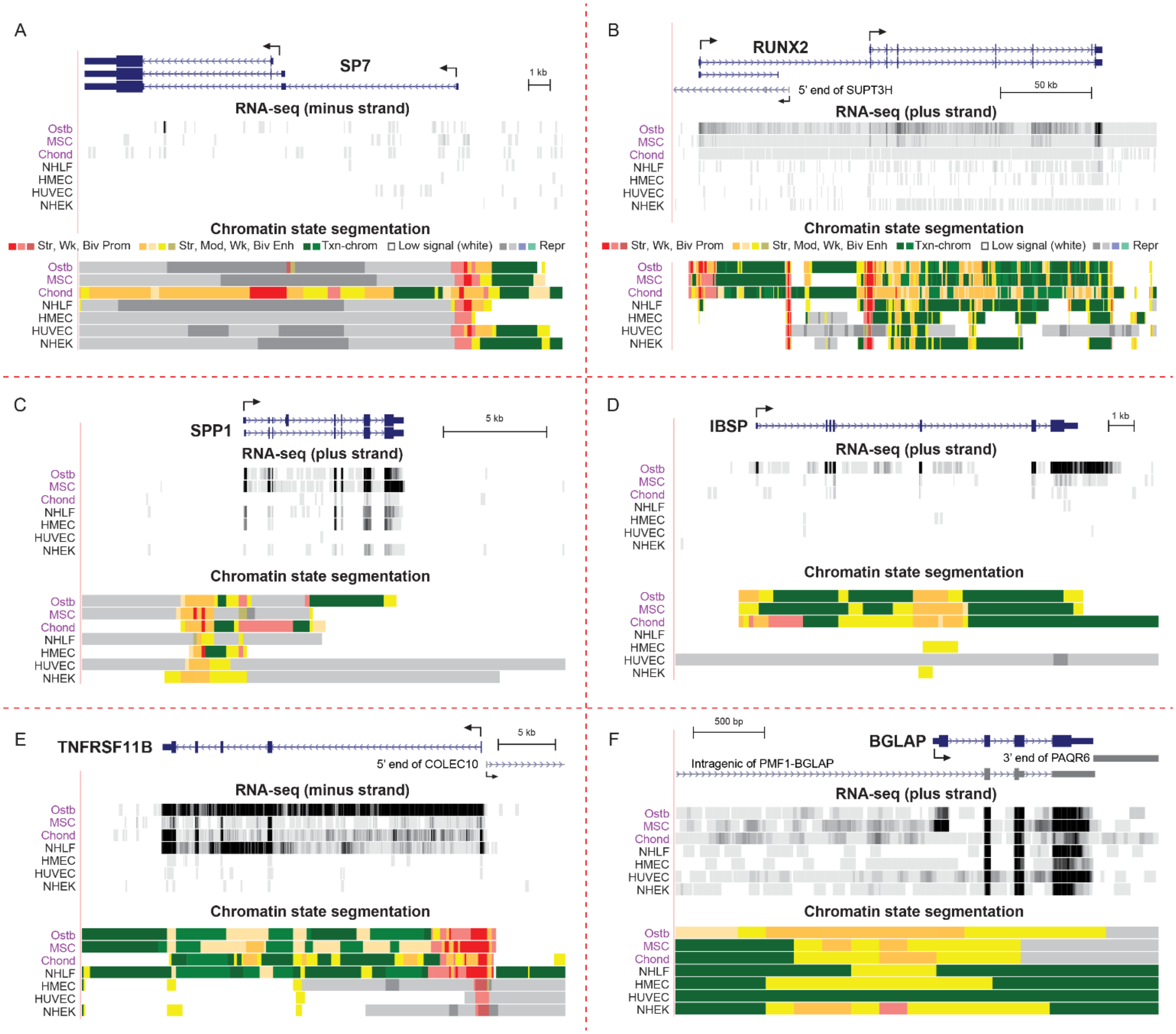


**Supplemental Fig. S1. Osteoblast primary cultures used for RNA-seq and epigenome profiling preferentially expressed five osteoblast-marker genes.** RNA-seq profiles (ENCODE)[1] and chromatin state segmentation profiles with the indicated color code (Roadmap)[3] for ostb markers were visualized in the UCSC genome browser (https://genome.ucsc.edu/). *(A) SP7/OSX*, a marker for pre-ostb,[5] was not expressed in these ostb cultures, indicating that they were not pre-ostb cultures. *(B-F) RUNX2* (expression starts when MSC commit to osteoblastogenesis and then increases),[6] *SPP1/OPN* (expressed in early and also later in ostb differentiation, http://biogps.org/#goto=genereport&id=20750), *IBSP* (a mid-to-late differentiation marker for ostb,[5] http://biogps.org/#goto=genereport&id=15891), *TNFRSF11B/OPG* (expressed in the early and middle stages of osteoblastogenesis, http://biogps.org/#goto=genereport&id=18383), and *BGLAP/OCN* (a terminal differentiation marker for ostb)[6] were all preferentially expressed in ostb and displayed chromatin segmentation profiles indicative of expression in these cells. With the exception of *BGLAP*, these ostb-associated genes have also been reported to be expressed in chond.[5, 7, 8] Ostb, osteoblasts; chond, chondrocytes; NHLF, post-natal lung fibroblasts (fib); HMEC, mammary epithelial cells; HUVEC, umbilical vein endothelial cells; NHEK, skin keratinocytes.

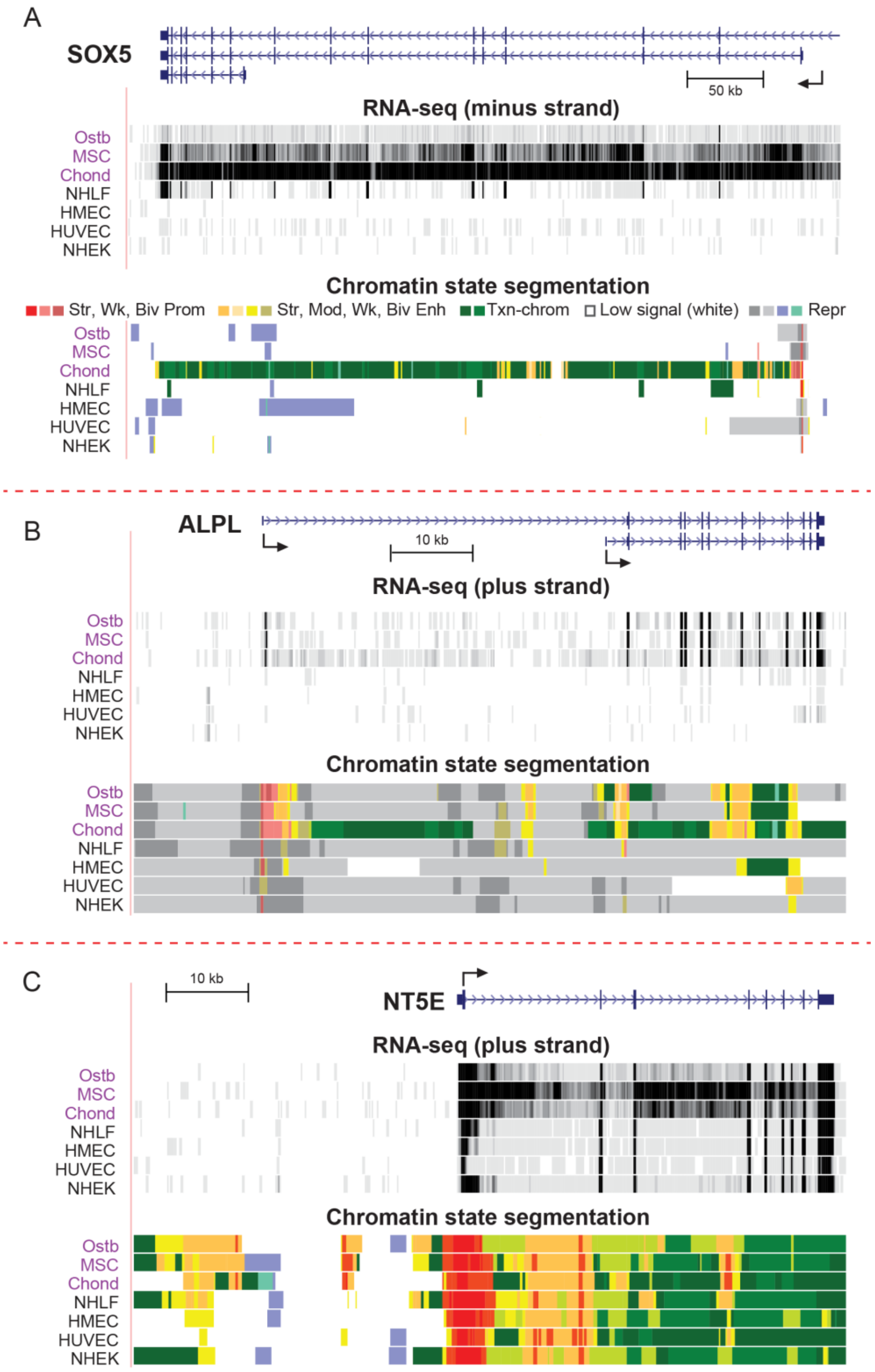


**Supplemental Fig. S2. Chondrocyte primary cultures used for RNA-seq and epigenome profiling selectively expressed the chondrocyte-marker gene *SOX5* and chondrocyte/osteoblast marker gene *ALPL*; bone marrow-derived mesenchymal stem cells over-expressed the corresponding marker gene *NT5E/CD73*.** *(A-C)* as in Supplemental Fig. S1. *SOX5*, chond marker gene.[9] *(B) ALPL*, chond and ostb marker[5, 10] displayed a chromatin segmentation profile indicative of expression most strongly in the chond sample, which had been derived *in vitro* from the MSC also used for epigenetic profiling. *(C)* MSC overexpressed *NT5E/CD73*, an MSC marker gene, relative to other cell types and, as expected, displayed negligible expression of the two negative markers for MSC (not shown, *CD5/PTPRC* and *CD34*).[11]

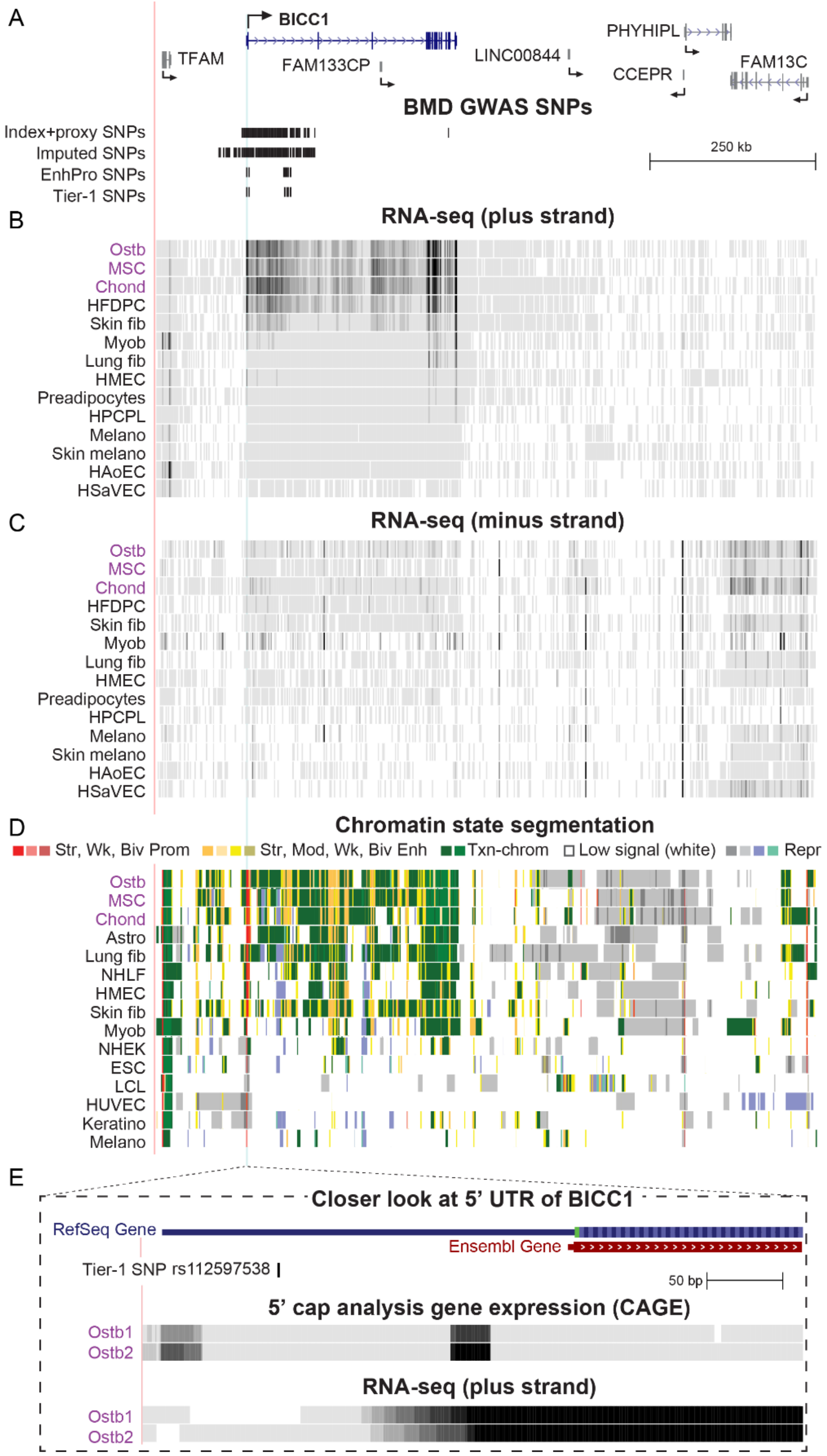


**Supplemental Fig. S3. The genes surrounding *BICC1* in a 1-Mb neighborhood have low expression in osteoblasts, unlike *BICC1*.** *(A)* The distribution of genes and several categories of BMD GWAS SNPs in the neighborhood of *BICC1* (chr10:60,135,702-61,136,560, hg19); EnhPro SNPs are SNPs overlapping strong enhancer (Enh) or promoter (Prom) chromatin preferentially in ostb; Tier-1 SNPs are the subset of EnhPro SNPs which have strong predictions of overlap of allele-specific transcription factor binding sites (TFBS) as described in the main text. *(B-C)* ENCODE strand-specific total RNA-seq (scale for the plus-strand, 0-300; for the minus-strand, 0-30). *(D)* Roadmap-derived chromatin state segmentation identifying strong (Str), moderate (Mod), weak (Wk), or bivalent (Biv; poised) Prom or Enh chromatin; chromatin with the H3K36me3 mark of actively transcribed regions (Txn-chrom), or repressed (Repr) chromatin. *(E)* Zoom-in to the 5' untranslated region (5' UTR) of *BICC1* (chr10:60,272,619-60,273,053) showing profiles from RNA-seq (scale, 0-1000) and CAGE (5'cap analysis gene expression; scale, 0-500) for mapping 5' caps of RNA; results from technical duplicates of an ostb sample are displayed. The RefSeq gene's 5' end was originally at the Ensembl gene's 5' end but recently was extended 0.2 kb further upstream, as shown. However, the CAGE and RNA-seq profiles indicate that the main end of this mRNA in ostb is ~65 bp upstream of the 5' end of the Ensembl gene structure. We assigned Tier-1 SNP rs112597538 as ~0.1 kb upstream of the *BICC1* TSS based on these CAGE and RNA-seq results. Ostb, MSC, chond, NHLF, HMEC, HUVEC, and NHEK (defined in the legend to Supplemental Fig. S1); HFDPC, follicle dermal papilla cells; fib, fibroblasts; myob, myoblasts; HPCPL, pericytes; melano, melanocytes from foreskin; HAoEC, aortic endothelial cells; HSaVEC, saphenous vein endothelial cells; astro, astrocytes; lung fib, fetal lung fib (IMR-90); ESC, embryonic stem cells; LCL, lymphoblastoid cell line; keratino, keratinocytes from foreskin.

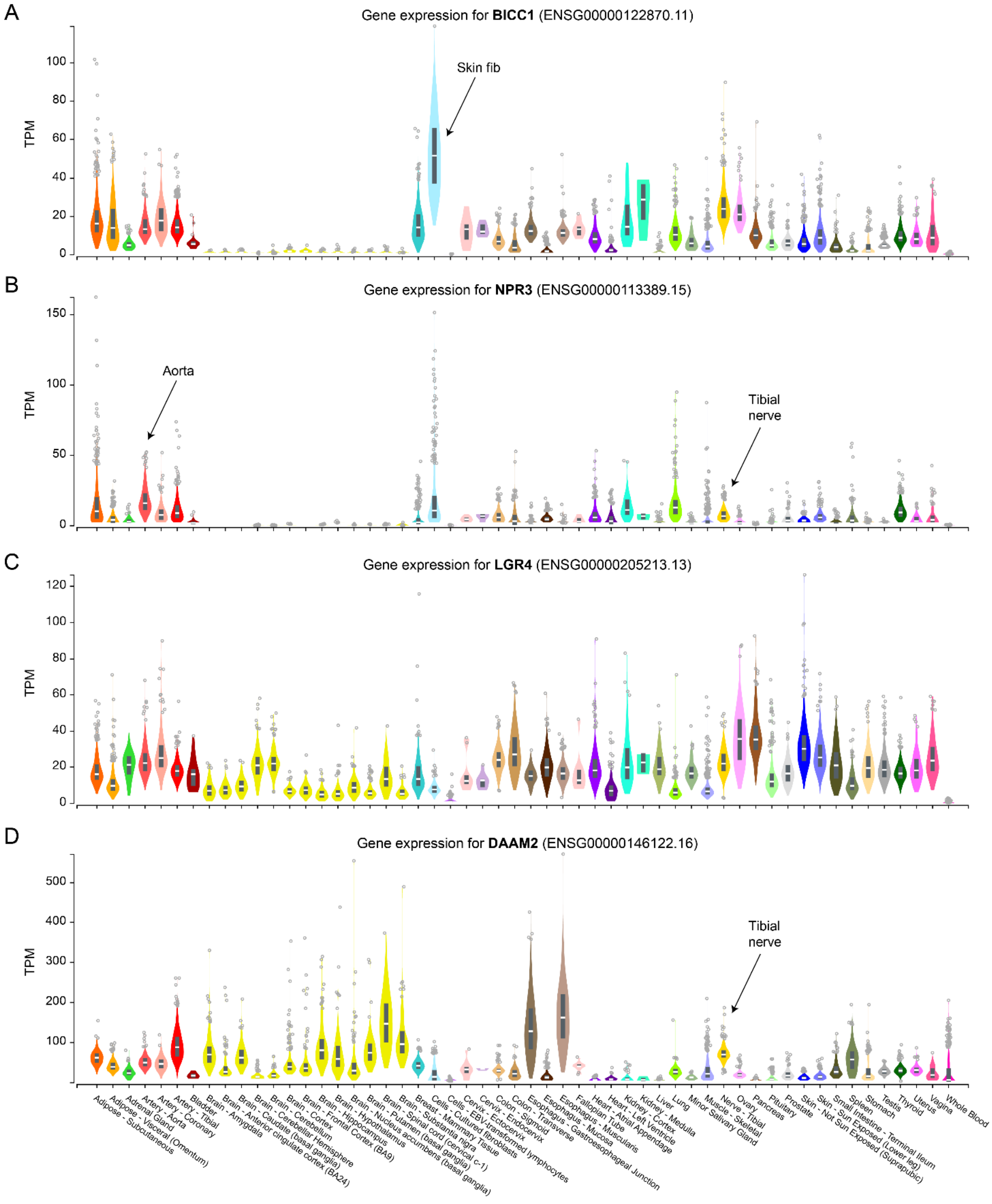


**Supplemental Fig. S4. Human tissue expression from GTEx for *BICC1, NPR3, LGR4,* and *DAAM2*.** *(A-D)* RNA-seq in transcripts per million (TPM) from GTEx[12] (https://www.gtexportal.org/). The median from many (usually hundreds) of samples is shown in violin plots for 51 human tissues and two types of cell cultures, primary human fibroblasts and B-cell lymphoblastoid cell lines.

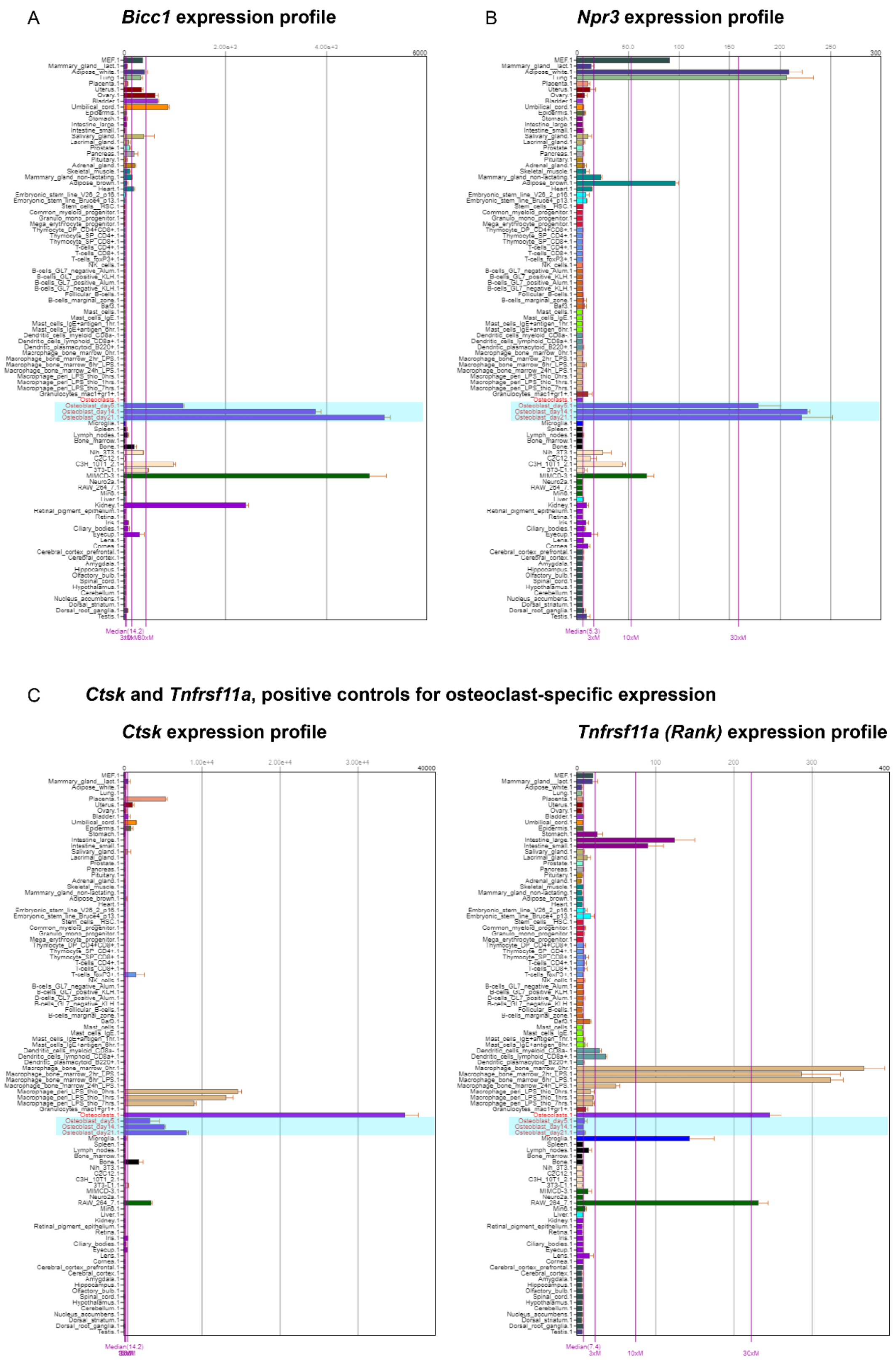


**Supplemental Fig. S5. Mouse expression microarray profiles from BioGPS for *Bicc1* and *Npr3*, as well as for the osteoclast- specific marker genes *Ctsk* and *Tnfrsf11a*.** *(A-C)* Blue highlighting, mouse ostb expression microarray profiles at day 5, 14 and 21 of differentiation (BioGPS,[13] http://biogps.org/). The expected osteoclast-specific expression of two osteoclast marker genes is seen in Panel C.

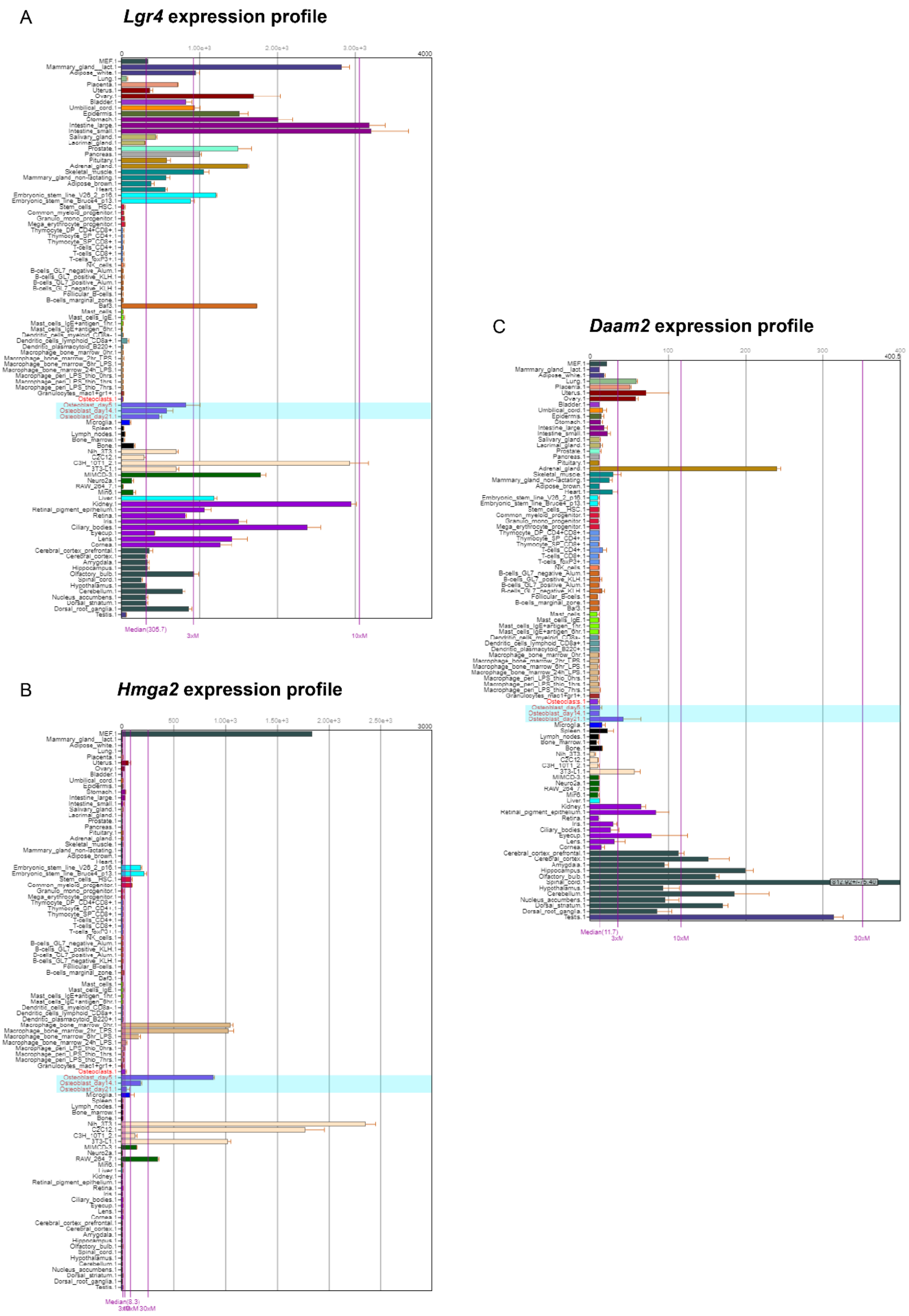


**Supplemental Fig. S6. Mouse expression microarray profiles from BioGPS for *Lgr4*, *Hmga2*, and *Daam2*.** (BioGPS,[13] http://biogps.org/).

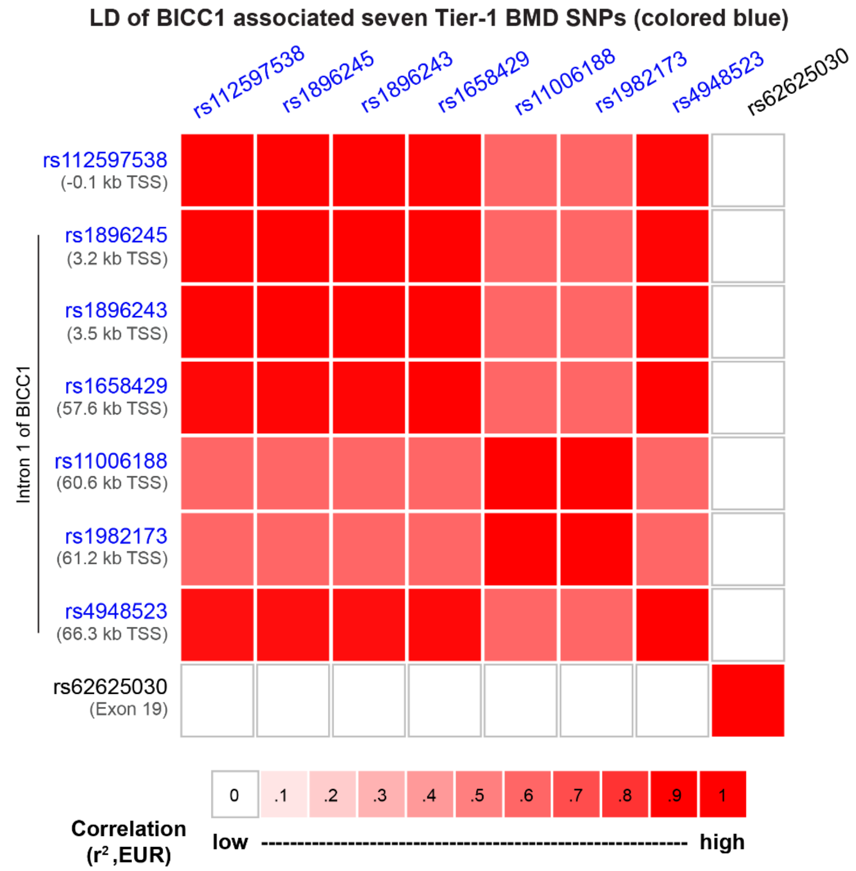


**Supplemental Fig. S7. Linkage disequilibrium (LD) matrix for seven *BICC1* associated Tier-1 BMD SNPs.** Seven Tier-1 SNPs for *BICC1* are in moderate to high LD ($r^2$ = 0.65 to 1, EUR) with each other but not with a *BICC1* missense variant rs62625030 identified by Morris et al.[14] (the point mutation at rs62625030 does not affect the structure of BICC1 as predicted by PSIPRED, http://bioinf.cs.ucl.ac.uk/introduction/)

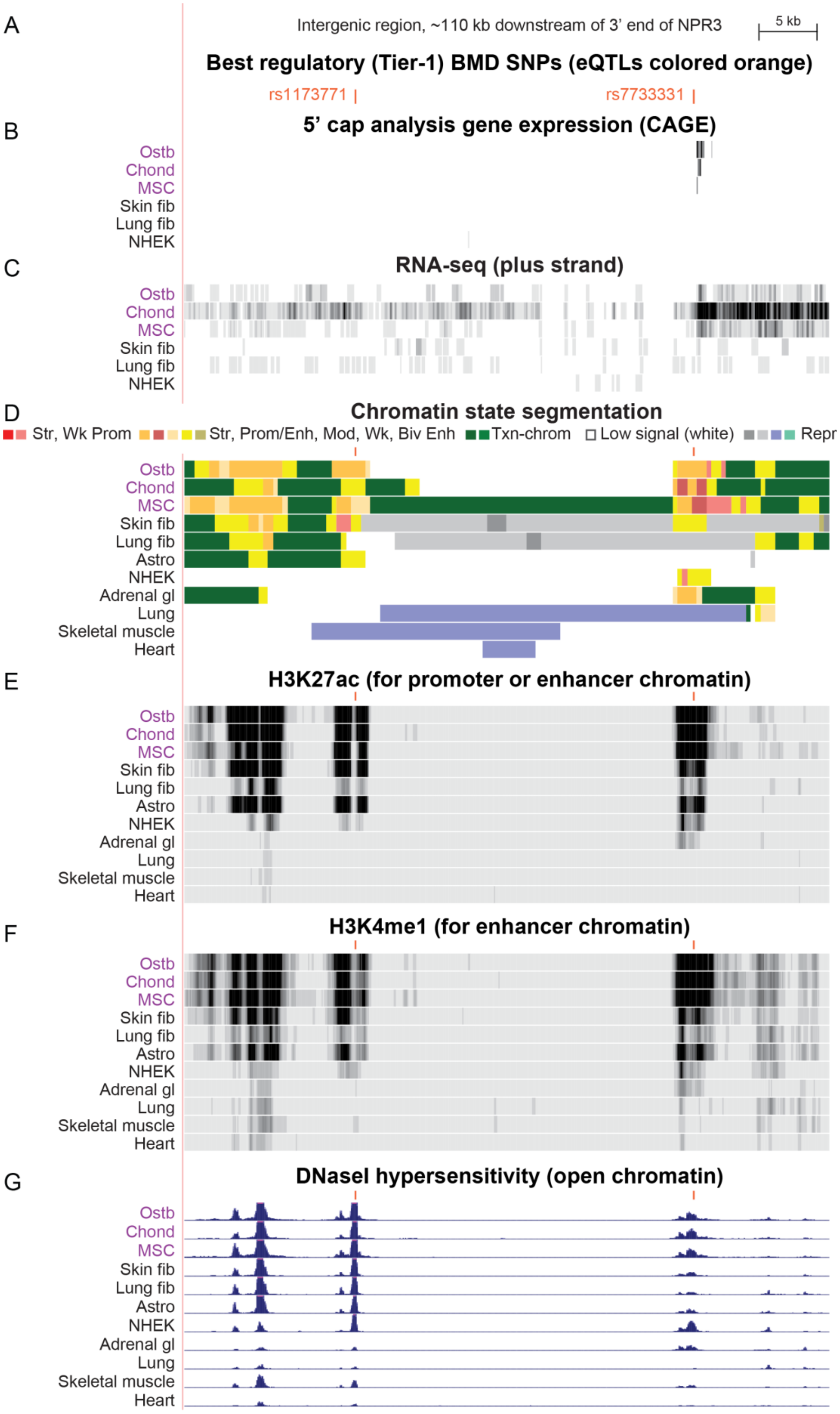


**Supplemental Fig. S8. Zoomed-in view of the intergenic region containing two *NPR3*-associated Tier-1 SNPs shows their proximity to the 5' end of the novel lincRNA.** *(A)* The two Tier-1 SNPs associated with *NPR3* (chr5:32,807,971-32,834,434). *(B)* Cell-specific CAGE signal from the plus-strand (scale, 0-5) as in Supplemental Fig. S3E. *(C)* Strand-specific total RNA-seq profile (scale, 0-30; in Fig. 3, the scale was 0-300, which obscured the lincRNA seen here). *(D)* Chromatin state segmentation; adrenal gl, adrenal gland. *(E)* H3K27ac signal; vertical viewing range 0-4. *(F)* H3K4me1 signal; vertical viewing range 0-4. *(G)* DNaseI hypersensitivity with the vertical viewing range set at 0-20.[3] Short orange bars above tracks in panels *(D-G)* are positions of Tier-1 SNPs.

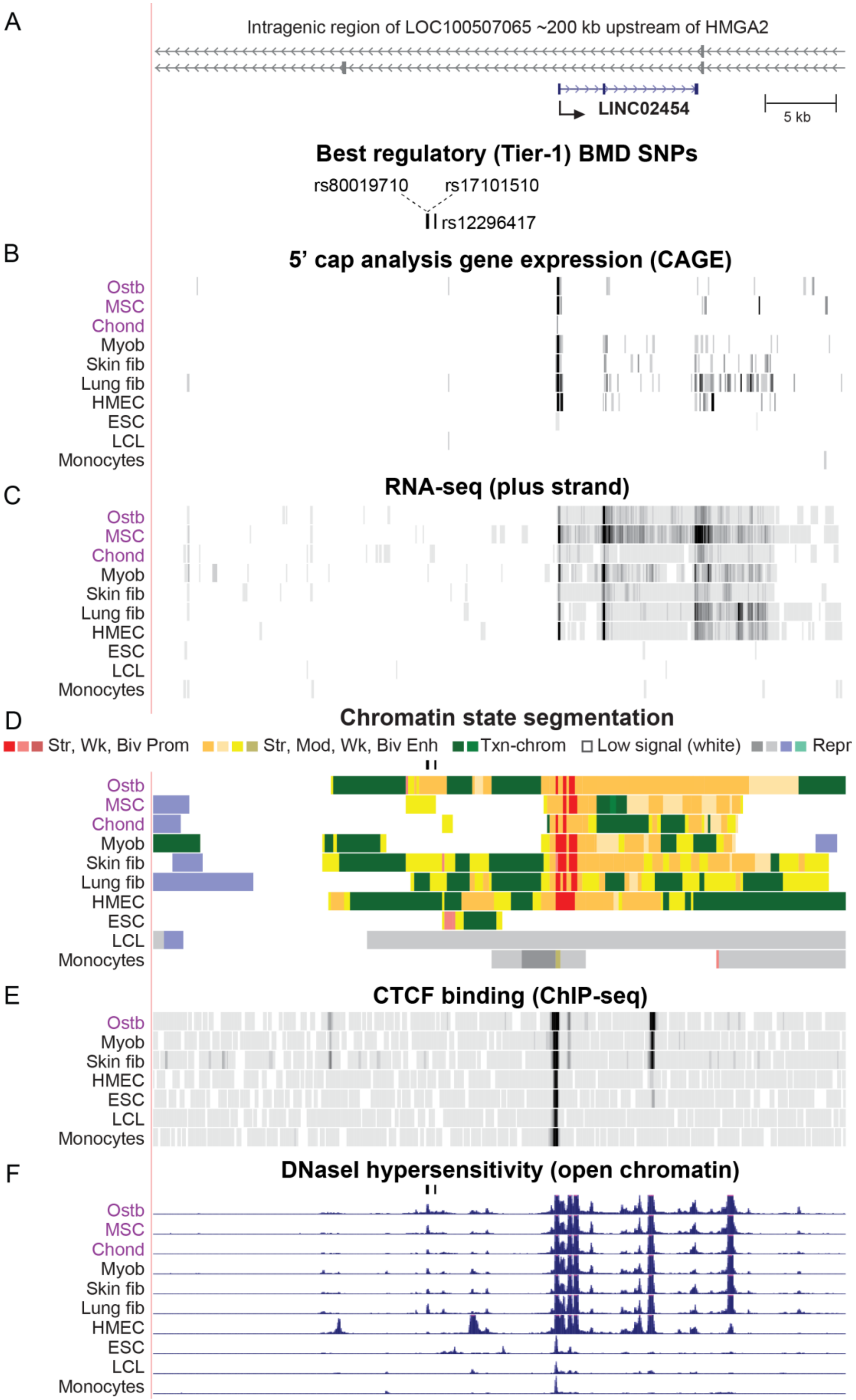


**Supplemental Fig. S9. Zoomed-in view of the *HMGA2* far-upstream region containing three Tier-1 SNPs shows the proximity of these SNPs to the 5' end of *LINC02454*.** *(A)* Tier-1 SNPs (chr12:65,967,381-66,017,380). *(B)* CAGE signal from the plus-strand as in Supplemental Fig. S3E (scale, 0-5). *(C)* Strand-specific total RNA-seq profile (scale, 0-90). *(D)* Roadmap-derived chromatin state segmentation. *(E)* CTCF binding determined by Roadmap ChIP-seq.[3] *(F)* DNaseI hypersensitivity, with the vertical viewing range set at 0-25.

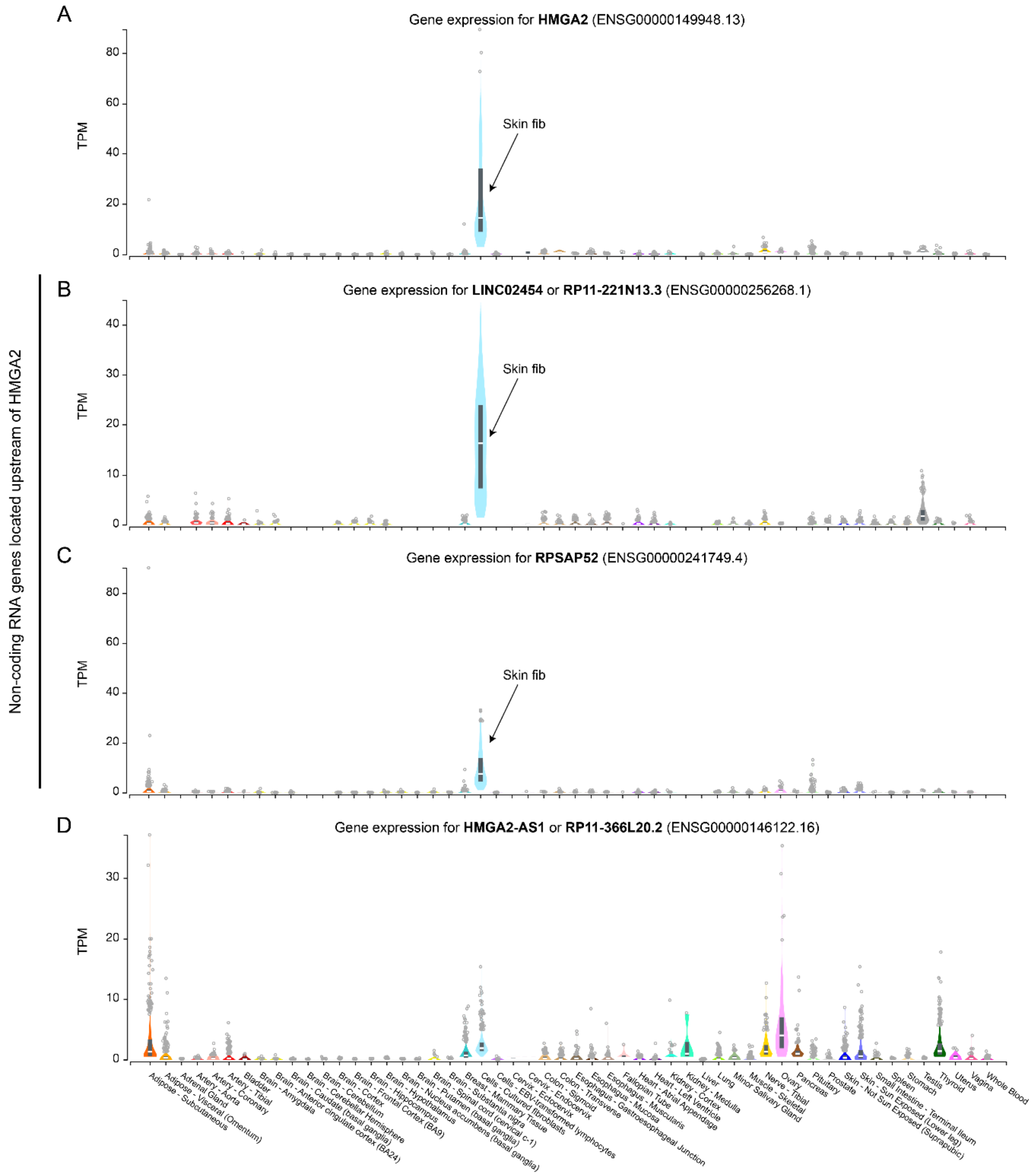


**Supplemental Fig. S10.** ***HMGA2*** **and two neighboring upstream non-coding RNA genes** ***RPSAP52,*** **and** ***LINC02454*** **have similar expression profiles while the** ***HMGA2-AS1*** **in the body of** ***HMGA2*** **has a different profile.** These profiles are derived from GTEx[12] (https://www.gtexportal.org/) and show median expression levels from many (usually hundreds) of tissue samples and two cell cultures (primary skin fibroblasts in light blue, which are next to B-cell lymphoblastoid cell lines, that had negligible expression).
